# Integrated genomics and transcriptomics reveal molecular insights into extreme dietary adaptation in vampire bats

**DOI:** 10.64898/2026.07.24.740446

**Authors:** Shenglin Liu, Mariella Bontempo Freitas, Sirlene Souza Rodrigues Sartori, Michele Albertini, Evgeny Leushkin, Ine Alvarez van Tussenbroek, Alejandro Gonzales-Irribarren, Ariadna E. Morales, Martin Pippel, Tom Brown, Alex Filipe Ramos de Sousa, Rosinéa Aparecida de Paula, Ilias Patmanidis, Willem Jespers, Leon Hilgers, Xueling Yi, Bernhard Bein, Yury Malovichko, Tilman Schell, Carola Greve, Sylke Winkler, Alexander Ben Hamadou, Moritz Blumer, Gisa Prange, Julio César Hechavarria Cueria, Manfred Kössl, York Winter, Sean Dilrosun, Stephert Daniël Bechan, Mark D. Engstrom, Deirdre Jafferally, Zacharias Norman, Francisco Sornoza, Liliana M. Dávalos, Burton K. Lim, Sonja C. Vernes, Michael Hiller

## Abstract

Vampire bats are the only tetrapods that feed exclusively on blood. To uncover the molecular changes associated with this extreme dietary specialization, we generated six new reference genomes, including genomes of all three vampire bat species, and integrated comparative analyses of gene sequence evolution (selection signatures, duplications, and losses) with transcriptomic data from six major organs to identify shifts in gene expression. Our integrative analyses reveal sequence or expression changes in 150 genes that provide insights into the molecular biology of sanguivory. Through comparative analyses and experiments, we show that the enlarged vampire bat stomach has increased connective tissue content, likely facilitating extreme expansion, is pH-neutral, and exhibits reduced mucus production, together providing molecular insights into its shift from a digestive to an absorptive organ for water, electrolytes, and vitamins. We further uncover pathway-level molecular changes associated with altered gastrointestinal motility; trypsin-dependent protein digestion; upregulated amino acid catabolism with key aspects diverging from other mammals; impaired dietary fat digestion counterbalanced by increased fatty acid synthesis; defective sugar metabolism and natural insulin deficiency; enhanced heme iron absorption; and adult splenic erythropoiesis. Together, these findings provide an integrated view of molecular changes associated with one of the most extreme dietary transitions among vertebrates.

## Introduction

Extreme dietary specializations impose strong and persistent selective pressures in nature, forcing organisms to reconcile severe nutritional imbalances with existing physiological and metabolic constraints. Deciphering how genomes adapt to such conditions reveals general principles of the evolutionary processes that remodel metabolism and physiology, with potential biomedical relevance (*1–6*). Obligate blood feeding (sanguivory), characterized by extreme nutrient bias, chronic caloric limitation, and metabolic states resembling features of human disorders such as insulin resistance, provides a powerful natural experiment for understanding the evolutionary mechanisms of extreme dietary adaptation.

Among more than 30,000 tetrapod species, obligate sanguivory has evolved only once and is restricted to three species of vampire bats (*7*, *8*): the common vampire bat *Desmodus rotundus*, the white-winged vampire bat *Diaemus youngii*, and the hairy-legged vampire bat *Diphylla ecaudata* (bat family Phyllostomidae, subfamily Desmodontinae). Vampire bats have long fascinated scientists for their extreme dietary specialization, which affects morphology, physiology, metabolism, and behavior.

Morphologically, these strictly nocturnal bats have well-developed terrestrial locomotion skills to approach prey, razor-sharp enamel-less upper incisors, salivary anti-coagulants to maintain blood flow, as well as advanced low-frequency hearing and infrared radiation sensing abilities (*9–11*). Their stomach transformed from a digestive to a distensible organ, used primarily for short term storage of large blood meals and rapid fluid absorption (*12*, *13*).

Physiologically, vampire bats have adapted to the high iron content and extreme nutrient bias of blood, which contains mostly protein (93%) and very little lipids (1%) and carbohydrates (1%) (*14*, *15*). To compensate for the low caloric content of their 78% water-containing diet, they consume large volumes in a single meal, often approaching half their body weight (*14*). Their low carbohydrate intake resulted in low basal insulin levels and a reduced glucose-stimulated insulin secretion response, resembling aspects of human type 2 diabetes (*16*). Limited glycogen and lipid stores make them very sensitive to starvation, with death occurring after 48–72 hours of fasting (*17*). Abundant dietary amino acids are their primary energy source and are rapidly used to fuel exercise such as running (*18*).

Behaviorally, vampire bats show exceptional social and cognitive adaptations. To mitigate risk of starvation, they regurgitate blood to roost mates that failed to forage, preferentially sharing with individuals recognized by social bonds formed through allogrooming and long-term social memory (*19–21*). They also adopt non-kin orphaned offspring and maintain long-lasting social bonds, even after captive bats are released into the wild (*22*, *23*). These advanced social behaviors are consistent with their large relative neocortical volume (*24*).

Several studies have shed light on genomic changes associated with sanguivory, revealing signatures of selection in genes involved in digestion, fat and vitamin metabolism, and immunity, a reduction of bitter taste receptor genes, and the loss of trehalase, the enzyme that digests the insect sugar trehalose (*25–30*). Comparative genomic studies also identified gene losses unique to the common vampire bat that relate to insulin secretion, glycogen storage, gastric function and their exceptional cognitive abilities (*31*).

However, previous studies have several limitations. First, almost all genomic studies focused on *Desmodus rotundus*, the only vampire bat species with a sequenced genome so far, leaving it unclear whether observed patterns are shared with *Diaemus youngii* or *Diphylla ecaudata*. Second, the lack of high-quality genomes of other phyllostomid bats limits taxonomic representation, with many studies often lacking the closest sister species of vampire bats. Third, previous work focused on single aspects of genomic change, such as positive selection or gene loss, leaving other types of molecular changes potentially important for adaptations, such as changes in gene expression, unexplored. Consequently, the molecular basis of sanguivory in vampire bats remains incompletely understood.

Here, we present high-quality genomes of six phyllostomid bats, including new haplotype-resolved assemblies of all three vampire bat species (for readability, we use the unique genus names *Desmodus*, *Diaemus* and *Diphylla* in the following). Our integrative analyses discovered changes in gene sequence and expression through comparative genomic and transcriptomics analyses of six major organs, providing molecular insights into their enlarged, non-acidic, absorptive stomach; altered gastrointestinal motility; trypsin-dependent protein digestion; upregulated amino acid catabolism; impaired digestion and increased *de novo* synthesis of fatty acids; defective sugar metabolism and natural insulin deficiency; enhanced heme iron absorption; and adult splenic erythropoiesis.

## Results

### Reference genomes of all three vampire bats

To sequence the *Diphylla* and *Diaemus* genome, we generated 42X and 39X coverage of PacBio HiFi reads and 61X and 66X Arima Hi-C data. For *Desmodus*, we generated 43 GB of additional HiFi data (increasing the total coverage to 57X) and re-assembled the new and existing read data (*31*). For all three species, we used Hifiasm in Hi-C mode and YaHS to generate haplotype-resolved chromosome-level assemblies (*32*, *33*), and performed dual haplotype curation to resolve remaining scaffolding and haplotype-sorting errors in the joint Hi-C map of both haplotypes. The haplotype assemblies of *Diphylla* and *Diaemus* have contig N50 values between 33 and 44 Mb, and at least 98.2% and 92.7% of these assemblies are contained in chromosome-level scaffolds (Fig. 1A-C, Supplementary Figure 1, Supplementary Table 1), matching the described karyotype of these species (*34*). Using the HiFi reads and Merqury (*35*), we estimate high base accuracies with QV values between 69 and 72, indicating only ∼1 error per 10 megabases. We note that these values represent an upper bound as the HiFi reads were used for assembly. In comparison to the previous *Desmodus* assembly, our new assembly improved both contig N50 (9 vs. 6.8 Mb and 11.3 vs. 8 Mb for haplotypes 1 and 2) and base accuracy (QV values 69 vs. 64 for both haplotypes).

**Figure 1:**
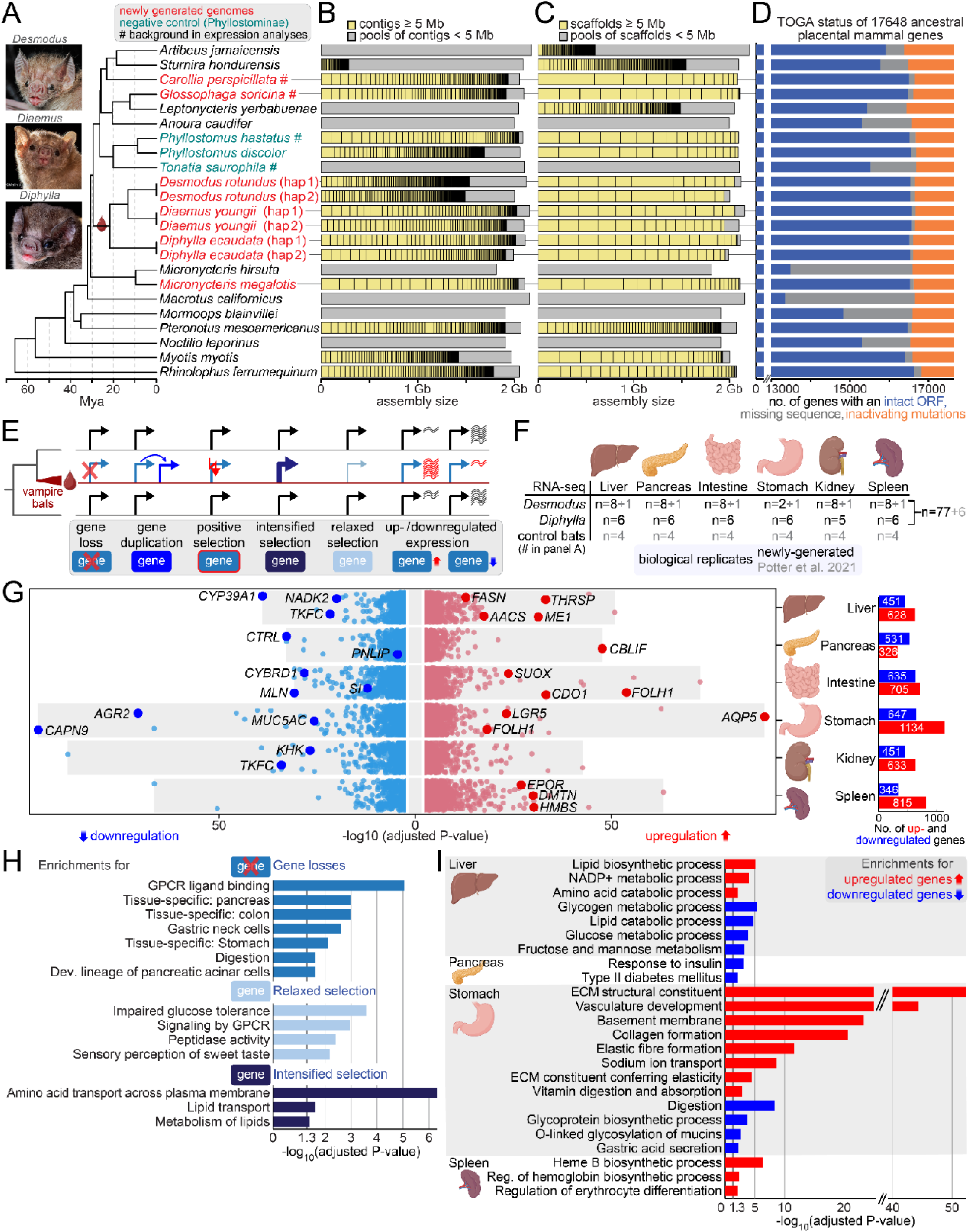
Comparative genomic and transcriptomic analyses illuminate extreme dietary adaptation in vampire bats. (A) Phylogeny of the bat species analyzed in this study. All nodes have 100% bootstrap support. Red indicates newly generated genomes, teal indicates the matched negative controls. Species used as background in the expression analyses are indicated by #. All 17 non-vampire species were used as background in the genomic screens. Insets show *Desmodus* (Photo credit Brock Fenton), *Diaemus* (©MerlinTuttle.org), and *Diphylla* (José G. Martínez-Fonseca). (B,C) Contig and scaffold size distributions, with boxes proportional to their size. (D) TOGA classification of ancestral placental mammal genes. (E) Illustration of comparative genomic and transcriptomic analyses to discover vampire bat changes in gene sequence and expression. (F) Newly-generated and existing organ transcriptomics data. (G) Significance (adjusted P-value scatter plots) and total number (bar charts) of differentially expressed genes in vampire bats relative to non-vampire species across six organs, highlighting some of the genes discussed in the text. (H,I) Significant enrichments of genes with sequence (H) and expression changes (I) associated with the discussed metabolic and physiological alterations in vampire bats. Supplementary Tables 11 and 12 list all enrichments.

Since assembly quality and completeness is important for comparative genomic analysis (*36*–*38*) and taxonomic coverage is key to understand which changes truly occurred in vampire bats, we generated three additional high-quality genomes of bats that cover other phyllostomid subfamilies, *Carollia perspicillata* (Carolliinae)*, Glossophaga soricina* (Glossophaginae), *Micronycteris megalotis* (Micronycterinae). For these species, we assembled primary genomes, having contig N50 values between 27 to 78 Mb and chromosome-level scaffolds (Fig. 1A-C, Supplementary Table 1).

To evaluate gene completeness, we used TOGA (Tool to infer Orthologs from Genome Alignments) (*39*) to classify 17,648 ancestral placental mammal coding genes into those that have an intact open reading frame, those with inactivating mutations (such as frameshifts, stop codon, exon deletions), and those with missing coding exons, often caused by assembly gaps. The new genomes contain between 16,486 and 16,570 intact genes (Fig. 1D, Supplementary Table 2), which is comparable or better than previous long and short read genomes (*26*, *28*, *40–43*). Applying compleasm (*44*) with the mammalia_ODB12 set of 12,277 near-universally conserved mammalian genes to our new genomes revealed a consistently high gene completeness ranging from 99.45% to 99.57% (Supplementary Table 1).

### Integrative analyses of genomic and transcriptomic changes associated with sanguivory

We compared the genomes of the three vampire bat species with those of 12 other phyllostomids and five non-phyllostomid bats. By including their closest sister and outgroup species, we were able to identify changes in the vampire bat lineage. To obtain a phylogeny of these bats, we used TOGA to identify 15,481 1:1 orthologous genes and used coalescence analysis (*45*, *46*) to infer a highly-supported, time-calibrated species tree (Fig. 1A, Supplementary Table 3).

Adaptations to sanguivory in vampire bats likely involve changes in both the sequence and expression of protein-coding genes. To identify gene sequence changes, we (i) used TOGA (*39*) to detect gene losses and duplications, (ii) applied the branch-site method aBSREL (*47*) to infer episodic positive selection, and (iii) used RELAX (*48*) to detect genes evolving under relaxed or intensified selective pressure to preserve the protein sequence (Fig. 1E, Supplementary Table 4). To uncover gene expression changes, we generated 77 RNA-seq datasets from six organs relevant for metabolism and physiology (stomach, small intestine, liver, pancreas, kidney, spleen) obtained from eight *Desmodus* and six *Diphylla* individuals (Fig. 1F, Supplementary Figure 2, Supplementary Table 5). We combined these datasets with available tissue-matched transcriptomes from one *Desmodus* and four non-vampire phyllostomid species (# in Fig. 1A) (*3*). Principal component analysis showed that newly-generated and published data clustered by tissue (Supplementary Figure 3), supporting organ-level expression coherence across species. We then used kb-python (*49*) to quantify gene expression and DESeq2 (*50*) to identify orthologous genes significantly up- or downregulated in one or both vampire bats, with many of these genes showing coherent expression shifts across both species (Fig. 1G, Supplementary Tables 6-8). These expression differences were overall robust to accounting for phylogenetic relationships among species (Supplementary Note 1, Supplementary Tables 9-10). Collectively, these integrative analyses provide a comprehensive view of gene sequence and expression changes in vampire bats.

Functional enrichment analyses highlight many metabolic and physiological processes that are altered in vampire bats (Fig. 1H,I, Supplementary Tables 11-14). Specifically, genes lost or evolving under relaxed selection are enriched for pancreas, colon, and stomach expression and functions related to digestion, taste perception, and glucose tolerance. Genes evolving under intensified selection are enriched in amino acid and lipid transport and lipid metabolism. Genes upregulated in the liver are enriched in amino acid catabolism and fatty acid synthesis, whereas downregulated genes are enriched in lipid and sugar catabolism. In the pancreas, downregulated genes are enriched in insulin response and type 2 diabetes–associated pathways. In the stomach, upregulated genes are enriched in extracellular matrix organization, vasculature development, and sodium and vitamin absorption (Supplementary Note 2), whereas downregulated genes are enriched in mucus production and gastric acid secretion. Finally, spleen-upregulated genes are enriched for erythropoiesis.

To assess whether the identified genes and functional enrichments are specific to vampire bats, we applied equivalent analyses to the Phyllostominae subfamily (teal in Fig. 1A), which comprises three omnivorous species as matched negative controls (*51*). We found virtually no overlap among the identified genes and limited overlap among the functional enrichments identified in the two lineages (Supplementary Note 3, Supplementary Tables 15-18), supporting the lineage specificity of many molecular signatures observed in vampire bats.

Guided by these functional enrichments and corroborated by new experiments, we describe in the following sections the potential contributions of 150 genes with sequence and/or expression changes in vampire bats (Supplementary Table 19) to their remodeled stomach, changes in gastrointestinal motility, adaptations in protein, fat, carbohydrate and iron metabolism, and splenic erythropoiesis. For all presented genes, we also show sequence alignments, amino acid and nucleotide substitution rates, and detailed gene expression profiles in Supplementary Data 1.

### Structural and size adaptations in the stomach

Vampire bats exhibit a markedly enlarged stomach (Fig. 2A). Among the most significantly upregulated genes in the vampire bat stomach are three stem cell marker genes, *LGR5*, *MSI1*, and *CD34* (rank 8, 17 and 24) (Fig. 2B). *LGR5* is expressed in mature pyloric glands and a subpopulation of chief cells, where it contributes to renewal of the gastric epithelium (*52*, *53*). *CD34*-expressing cells promote the maintenance of *LGR5*-expressing intestinal epithelial stem cells (*54*). *MSI1*, a gastric stem cell marker, controls the switch of intestinal stem cells from a quiescent to an active state and is required for regeneration of gastric mucosal epithelium upon injury (*55*, *56*). Upregulation of these genes may contribute to the distinct cell-type composition of the vampire bat stomach that has acinar rather than deep tubular gastric glands and fewer parietal and chief cells (*57*) (Supplementary Figure 4). Furthermore, since all three genes promote maintenance of stem cell identity, their upregulation may also contribute to stomach enlargement.

**Figure 2:**
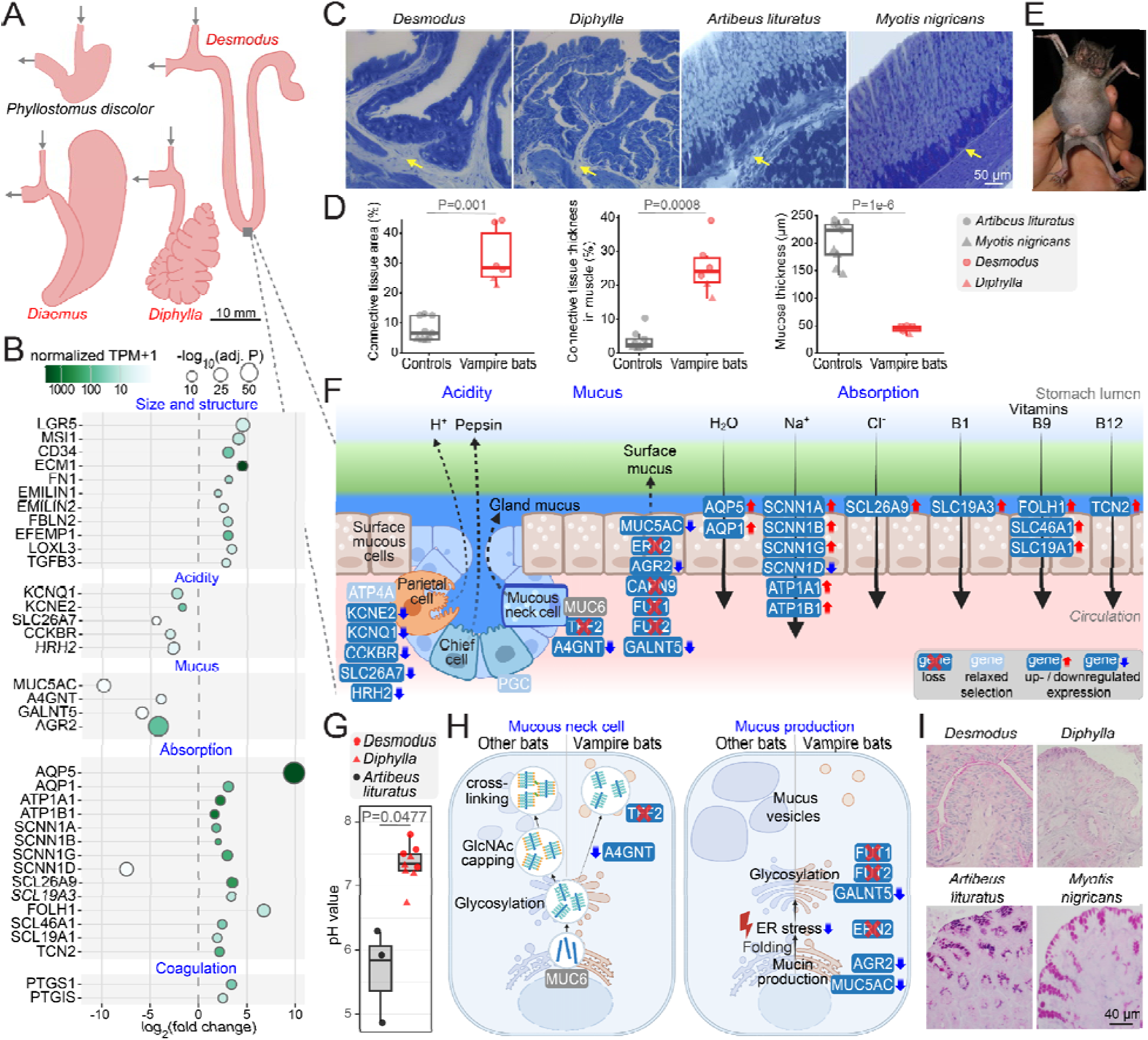
Molecular mechanisms involved in the enlarged, non-acidic, and absorptive vampire bat stomach. (A) Stomach size and morphology, drawn to scale based on (*65*, *66*), comparing the omnivorous *Phyllostomus discolor* with vampire bats. Inward arrows indicate esophagus, outward arrows duodenum. (B) Differential gene expression in the stomach, showing fold changes relative to non-vampire expression-background species (# in Fig. 1A; X-axis), expression levels (normalized TPM, color gradient) and significance (circle size). Genes are ordered as discussed in the text. (C) Histology (Toluidine blue staining) of stomach walls. Yellow arrows highlight connective tissue content. Scale bar is 50 µm for all images. (D) Quantification of connective tissue content and mucosa thickness, assessing differences with two-tailed Welch’s t-tests. (E) Fed *Desmodus* individual, illustrating the expanded blood-filled stomach (photo credit Jon Flanders). (F) Illustration of sequence and expression changes of genes involved in reduced gastric acidity, mucus production and increased absorption. (G) pH meter measurements show that vampire bats have a significantly higher stomach pH that is near-neutral. Two-tailed Welch’s t-test was used. (H) Illustration of gene changes relevant for mucus neck cells (left) and general mucus production (right). (I) Histological analysis with Periodic acid-Schiff/Alcian blue staining shows reduced gastric mucus in vampire bats.

Upregulated genes in the vampire bat stomach are strongly enriched for functions related to “extracellular matrix (ECM) structural constituent”, “basement membrane”, “collagen formation”, and “elastic fiber formation” (Fig. 1I). Many of these genes are normally expressed in connective tissue: *ECM1* acts as “biological glue” for various ECM and basement membrane components (*58*), fibronectin (*FN1*) forms fibrillar networks crucial for cell adhesion and ECM organization (*59*), elastic fiber components (*EMILIN1*, *EMILIN2*) contribute to tissue elasticity (*60*), fibulins (*FBLN2*, *EFEMP1*) associate with elastic fibers and stabilize basement membranes (*61*, *62*), *LOXL3* enhances ECM stiffness by crosslinking collagen and elastin (*63*), *TGFB3* regulates ECM and cellular adhesion components and contributes to ECM stiffness (*64*), and numerous collagen genes (Fig. 2B).

To investigate the basis of these gene upregulations, we performed histological comparisons of the stomachs of *Desmodus*, *Diphylla*, the fruit-eating phyllostomid bat *Artibeus lituratus*, and the insect-feeding bat *Myotis nigricans*. Image analysis and quantification revealed that vampire bats have significantly increased connective tissue content, including within the muscular layer, and reduced mucosal thickness (Fig. 2C-D, Supplementary Figure 5, Supplementary Table 20). This marked increase in connective tissue likely provides the mechanical support needed for the extreme stomach expansion that occurs when vampire bats ingest blood volumes approaching 60% of their body weight (*15*) (Fig. 2E). Together, our analyses provide molecular insights into structure and size adaptations of the vampire bat stomach.

### Non-acidic stomach

In most mammals, the stomach is primarily an acidic digestive organ. We found that *ATP4A*, the catalytic subunit of the gastric H^+^/K^+^-ATPase proton pump responsible for gastric acid secretion (*67*), evolves under relaxed selection in vampire bats (Fig. 2F). Furthermore, genes downregulated in the vampire bat stomach are enriched for “gastric acid secretion” (Fig. 1I), including those encoding the K⁺ channel KCNQ1 and its regulatory subunit KCNE2, which replenish luminal K⁺ (*68*), the histamine-stimulated chloride/bicarbonate exchanger SLC26A7, which mediates parietal cell chloride uptake (*69*), and the parietal cell receptors for the gastric acid secretagogues gastrin (CCKBR) and histamine (HRH2) (*70*) (Fig. 2B,F). Inhibition or knockout of all these genes impairs gastric acid secretion, and *CCKBR* knockout reduces parietal cell numbers (*71–75*), consistent with previous anatomical studies reporting fewer parietal cells without canaliculi and fewer gastrin-producing cells (*12*, *76*, *77*).

Together, these genomic and transcriptomic signatures suggest reduced stomach acidity in vampire bats. To test this prediction, we measured stomach acidity in *Desmodus* (n=4), *Diphylla* (n=5) and, as a control, in the phyllostomid bat *Artibeus lituratus* (n=3). Indeed, vampire bats have a near-neutral pH (mean 7.36 ± 0.29), while *Artibeus* has a weakly acidic stomach (mean 5.67 ± 0.71, two-tailed Welch’s t-test P=0.048; Fig. 2G, Supplementary Table 21).

Three factors likely underlie the non-acidic stomach. First, because blood has a high buffering capacity, acidifying the enlarged vampire bat stomach would require substantial acid production, which is likely energetically prohibitive. Second, the altered gastric mucus profile (below) likely makes the stomach vulnerable to acid-induced damage. Third, and probably most important for vampire bats, their stomach shifted from a digestive to an absorptive organ (below).

### Remodeled gastric mucus profile

Gastric mucus protects the stomach epithelium. Our integrative analysis revealed losses and downregulation of several genes related to mucus composition and function (Fig. 2B,F,H), supported by the enrichment of stomach-downregulated genes in mucin production and glycosylation (Fig. 1I). *MUC5AC*, the most abundant mucin in a typical mammalian stomach (*78*), is not expressed in the vampire bat stomach. The other major gastric mucin, *MUC6*, is robustly expressed; however, its co-expressed binding lectin *TFF2* (*79*, *80*) is lost in vampire bats (Supplementary Figure 6). Interactions between TFF2 and MUC6 are required to stabilize the inner mucus layer and *TFF2*-deficient mice exhibit reduced mucus viscoelasticity and mucosal thickness (*81*, *82*). TFF2 binds specifically to an α1,4GlcNAc-capped glycan on MUC6 (*79*). Consistent with *TFF2* loss, the α1,4GlcNAc-capping enzyme *A4GNT* is downregulated in the vampire bat stomach.

Several other genes normally expressed in gastric mucus-producing cells (*83*) show evidence of loss, relaxed selection, or downregulation in the vampire bat stomach (Fig. 2B,F,H, Supplementary Figures 7-11, Supplementary Notes 4-5). These include the protease *CTSE* (*84*), the endocytosis regulator *REP15* (*85*, *86*), the gastric mucosal defense factor *CAPN9* (*87*), the mucus glycosylation enzymes *FUT1*, *FUT2* and *GALNT5* (*88–91*), and the disulfide isomerase *AGR2*, which is essential for mucus production (*92*). Furthermore, vampire bats lost *ERN2*, an endoplasmic reticulum (ER) stress sensor that regulates the unfolded protein response factor XBP1 (*93*), potentially reflecting reduced ER stress due to diminished production of large glycoproteins, such as the >3500 amino acid mammalian MUC5AC.

Changes and losses of these genes predict reduced gastric mucus production. To experimentally investigate this, we performed histological analyses of the fundic stomach region in *Desmodus*, *Diphylla*, *Artibeus lituratus*, and *Myotis nigricans*. Periodic acid-Schiff/Alcian blue staining, which detects mucus and mucus-secreting cells, showed abundant staining in *A. lituratus* and *M. nigricans*, but markedly reduced staining in *Desmodus* and *Diphylla* (Fig. 2I).

Several factors likely account for the altered gastric mucus profile in vampire bats. First, the protective function of mucus against abrasive food and gastric acid is largely obsolete in species that consume only liquid food and lack an acidic stomach. Second, mucin synthesis is energetically costly (*94*), which may contribute to the absence of *MUC5AC* expression. Third, because vampire bats lack MUC5AC and the mucus-crosslinking factor TFF2, which form a stiff, highly branched, viscoelastic mucus network (*80*, *82*, *95*) (Fig. 2H), their gastric mucus likely has reduced viscoelasticity. This reduction may facilitate extreme stomach distensibility (Fig. 2E) and may contribute to their unique ability to absorb water in the stomach (below). Hence, remodeling the gastric mucus profile is likely adaptive from an evolutionary perspective.

### Functional shift of the stomach from digestion to absorption

Unlike other mammals, the highly-vascularized stomach of vampire bats can efficiently absorb water, which facilitates rapid water extraction and excretion (*96–98*); however, the underlying molecular mechanisms remained unclear. Our transcriptome data suggest that vampire bats primarily utilize aquaporin AQP5, a water channel normally expressed mainly in salivary, lacrimal and sweat glands, and the respiratory tract (*99*, *100*), for water absorption in the stomach (Fig. 2B,F). Whereas *AQP5* expression is very low in the stomach of non-vampire bats (average rank 10362), it is the top upregulated gene in the vampire bat stomach (943-fold increase) and ranks as the 23rd most highly expressed gene in this organ. Another water channel, *AQP1*, is 9-fold upregulated in the vampire bat stomach. Since expression of *AQP5* or *AQP1* induces cellular water permeability (*101*, *102*), the gain of expression of these water channels provides a plausible mechanism for water absorption in the vampire bat stomach.

We further discovered additional gene expression changes that support a transformation of the vampire bat stomach from a digestive to an absorptive organ for electrolytes. Blood has an exceptionally high sodium content, ∼34-fold higher than a typical human diet (Supplementary Note 6), and common vampire bats excrete most dietary sodium through urine within the first hour after feeding (*103*). This rapid excretion indicates that dietary sodium is already absorbed in the stomach along with water. Indeed, consistent with the enriched term “sodium ion transport” (Fig. 1I), our transcriptome data show that both subunits of the sodium/potassium ATPase (*ATP1A1*, *ATP1B1*) and all three subunits of the epithelial sodium channel αβγ-ENaC (*SCNN1A*, *SCNN1B*, *SCNN1G*), which are normally important for intestinal absorption and renal reabsorption of sodium (*104*), are highly upregulated in the vampire bat stomach (Fig. 2B,F). Interestingly, the δ-ENaC subunit (*SCNN1D*), which can replace the α subunit to form δβγ-ENaC, is downregulated in the stomach. This is consistent with the pH-neutral environment in the vampire bat stomach: δβγ-ENaC is inactive at pH>6.5, whereas αβγ-ENaC exhibits maximal activity under pH-neutral conditions (*105*, *106*). High *ATP1A1*/*ATP1B1* expression further aligns with previous observations of a high sodium/potassium ATPase activity in the common vampire bat, and sodium absorption could create an osmotic drive that enhances the water absorption (*96*). High rates of sodium cation uptake raise the question of how vampire bats maintain electroneutrality. We found the chloride transporter gene *SLC26A9* (*107*, *108*) to be 10-fold upregulated and among the most upregulated genes in the vampire bat stomach (Fig. 2B,F); this upregulation may enhance chloride anion absorption to facilitate electroneutrality.

In addition to electrolytes, we detected expression shifts suggesting uptake of the essential vitamins B1, B9, and B12 in the stomach (Fig. 2B,F), consistent with the enrichment of “vitamin digestion and absorption” (Fig. 1I). Intestinal uptake of vitamin B1 (thiamine) is mediated by *SLC19A3* (*109*, *110*). We found this gene to be upregulated in the vampire bat stomach, suggesting that thiamine uptake also occurs in this organ. Intestinal uptake of vitamin B9 (folate) requires FOLH1, which hydrolyzes the dietary form of folate (polyglutamate-folate), enabling monoglutamate-folate uptake across the brush border membrane by SLC46A1 and SLC19A1 (*111*, *112*). All three genes are upregulated in the vampire bat stomach. These genes are also highly expressed in the intestine, with *FOLH1* being the third-most strongly upregulated gene (602-fold) in this organ. Finally, *TCN2*, encoding the vitamin B12 transport protein transcobalamin II (*113*), is upregulated in the stomach. Together, these findings indicate enhanced folate and vitamin B12 absorption in vampire bats.

Finally, *PTGS1* and *PTGIS*, encoding the two enzymes that produce platelet-inhibiting prostacyclins, are upregulated in the vampire bat stomach (Fig. 2B), potentially helping to limit clotting of ingested blood and facilitating its regurgitation to roost mates. Overall, these expression shifts provide strong evidence that the vampire bat stomach has been evolutionarily repurposed into an absorptive organ.

### Altered gastrointestinal motility

We identified losses of several genes regulating gastrointestinal motility in vampire bats (Supplementary Figures 12-13). The gene encoding the peptide hormone motilin (*MLN)* is lost in *Diaemus* and *Diphylla*, and the motilin receptor gene *MLNR* is lost in *Desmodus* and *Diaemus* and exhibits a strong signal of relaxed selection in vampire bats. In most mammals, motilin regulates fasting state gastrointestinal motility by initiating phase III contractions of the migrating motor complex, which clear the intestinal lumen during the interdigestive period (*114*). Consistent with losses of *MLN* and *MLNR*, motilin-producing endocrine cells are absent from the gastrointestinal tract of *Desmodus* (*115*), suggesting that intestinal cleansing is less important for species adapted to an exclusively liquid blood diet with little indigestible material. In addition, the galanin receptor *GALR2*, which stimulates gastrointestinal motility (*116*, *117*), is lost in all vampire bats. Together, these gene losses indicate altered gastrointestinal motility in vampire bats.

To experimentally investigate this, we examined gastrointestinal movement in *Desmodus* (n=3) and *Diphylla* (n=2), alongside three control species. In contrast to *Phyllostomus discolor* (n=2), *Artibeus lituratus* (n=2), and mice (n=3), vampire bats exhibited no detectable gastrointestinal motility (Supplementary Videos 1-5). We deliberately stimulated motility using neostigmine (0.5 mg/mL), a cholinesterase inhibitor that induces gastrointestinal motility in mammals. Whereas neostigmine elicited the expected robust movements in *Artibeus lituratus*, *Phyllostomus discolor*, and mice, vampire bats showed no detectable response (Supplementary Videos 1-5). Together, these experiments reveal altered gastrointestinal motility in vampire bats, consistent with their exclusively liquid diet.

### Reduction of protein digestive enzymes and increased trypsin dependence

Our analyses uncovered losses or relaxed evolutionary constraint on several genes encoding zymogens involved in digesting dietary proteins (Fig. 3A, Supplementary Figures 14-17). *PGC,* encoding the inactive pepsinogen C precursor secreted by gastric chief cells, evolved under relaxation in vampire bats (Fig. 2F). Since pepsinogen C requires an acidic environment (pH<5) for autocatalytic activation (*118*), the near-neutral stomach pH in vampire bats provides a potential explanation for relaxation. Among the digestive enzymes secreted by the exocrine pancreas, chymotrypsinogen B2 (*CTRB2)* and the chymotrypsin-like protease (*CTRL)* are lost in all vampire bats. Elastase-1 (*CELA1*) is lost in *Diaemus*, evolves under relaxation in all vampire bats, and is not expressed in the *Desmodus* pancreas (Supplementary Table 7). Pancreatic carboxypeptidase A2 (*CPA2*) is lost in *Diphylla* only. Although the reduced repertoire of protein-digestive enzymes in vampire bats may seem counterintuitive, blood provides a limited diversity of readily digestible proteins, dominated by the soluble globular proteins hemoglobin and albumin. Hence, the larger enzyme repertoire required to digest the protein complexity contained in other diets may be less important for vampire bats.

**Figure 3:**
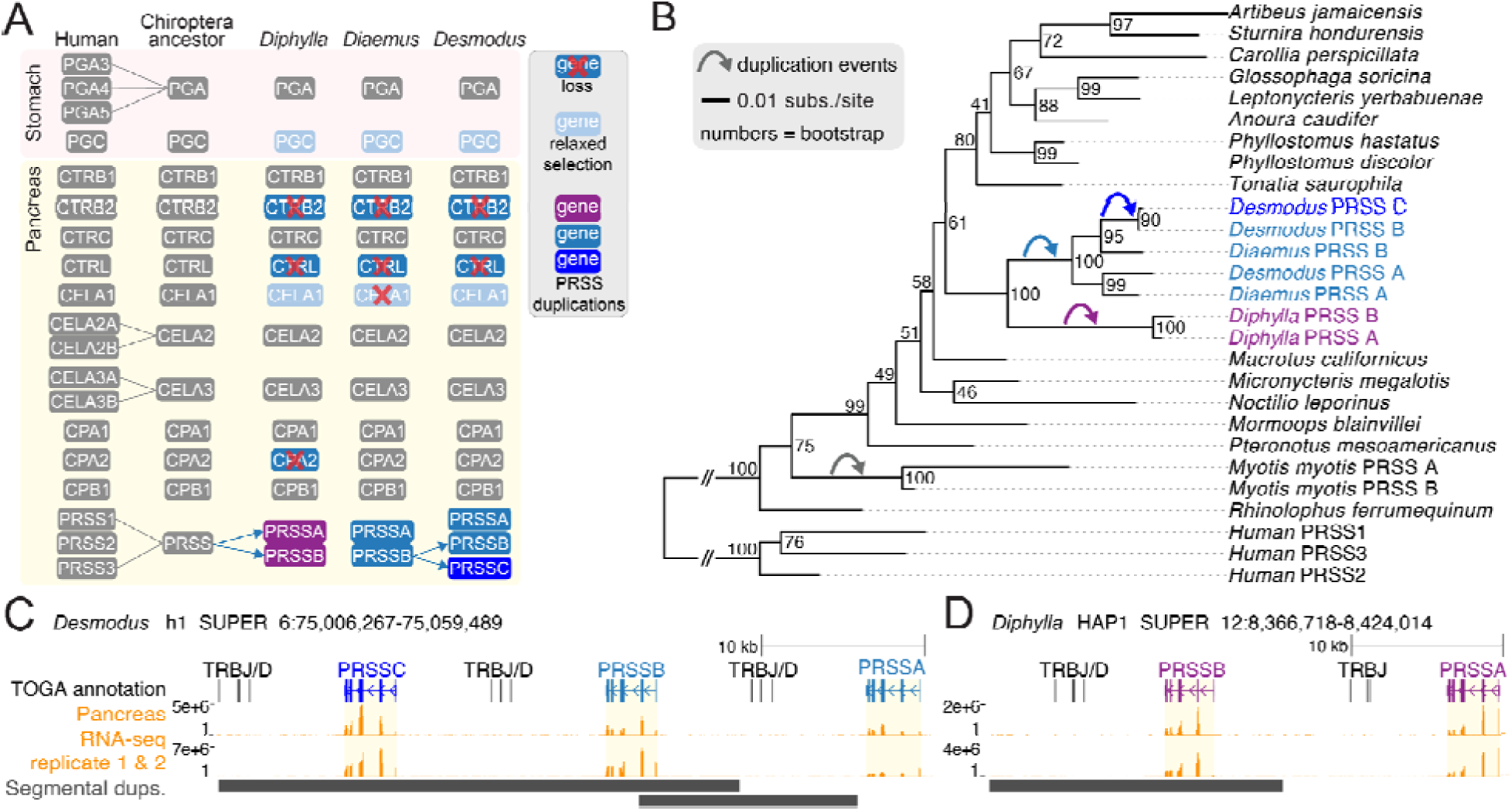
Protein digestion and independent trypsin duplications. (A) Evolution and changes in protein-digestive enzymes in the stomach and pancreas. Trypsin (PRSS) gene colors match panels B-D. Genes in grey background show no detected changes. (B) Maximum-likelihood tree generated with IQ-TREE v1.6.12 (*122*) showing that the phyllostomid ancestor possessed a single pancreatic trypsin gene that is co-orthologous to human *PRSS1-3*. This gene independently duplicated in vampire bats (curved arrows). *Myotis myotis* also exhibits an independent trypsin duplication, supported by its sister species *Myotis daubentonii* (not shown). (C,D) Genome browser screenshots showing the expanded trypsin genes in *Desmodus* (C) and *Diphylla* (D) (top) and that they are all expressed in pancreas. Segmentally duplicated genomic regions (black bars, bottom) were inferred with BISER (*123*).

In contrast to losses of digestive proteases, we found that the pancreatic trypsin-encoding gene *PRSS1* expanded into two copies in *Diphylla* and *Diaemus* and three copies in *Desmodus*. Interestingly, gene tree analysis revealed a complex duplication history with independent trypsin duplications in the *Desmodus*/*Diaemus* and in the *Diphylla* lineage, followed by another duplication in *Desmodus* (Fig. 3B, Supplementary Figure 18). All trypsin copies are expressed in the pancreas of *Diphylla* and *Desmodus* (Fig. 3C,D), likely increasing the trypsin pool. Furthermore, because CTRB2 and CTRL can cleave and degrade trypsinogens (*119*, *120*), their loss could also contribute to an increased trypsin pool. Finally, *SPINK1*, a serine protease inhibitor that protects the pancreas from prematurely activated trypsinogen (*121*), is upregulated in the vampire bat pancreas (Supplementary Table 6). Overall, our analysis indicates that, while the importance of several digestive proteases has decreased, trypsin may have become increasingly important for vampire bats, likely reflecting the low diversity and high digestibility of blood proteins.

### Increased reliance on intestinal absorption and renal reabsorption of amino acids

After protein digestion, amino acids are absorbed in the small intestine by multiple transporters (*124*). In vampire bats, many intestinal amino acid transporters show slightly elevated expression, with *SLC6A20* and *SLC16A10* significantly upregulated, and multiple transporters evolve under intensified selection (Fig. 4A). Notably, *SLC6A19*, encoding the only neutral amino acid transporter at the apical membrane of enterocytes, shows strong intensified selection and is the most highly expressed amino acid transporter in the small intestine of bats. *SLC6A19* is essential for intestinal absorption of branched-chain and aromatic amino acids, which is severely impaired in humans or mice lacking this gene (*124*, *125*). In the large intestine, two additional transporters for neutral amino acids (*SLC1A5*) and glycine, proline, alanine (*SLC36A1*) are also under intensified selection.

**Figure 4:**
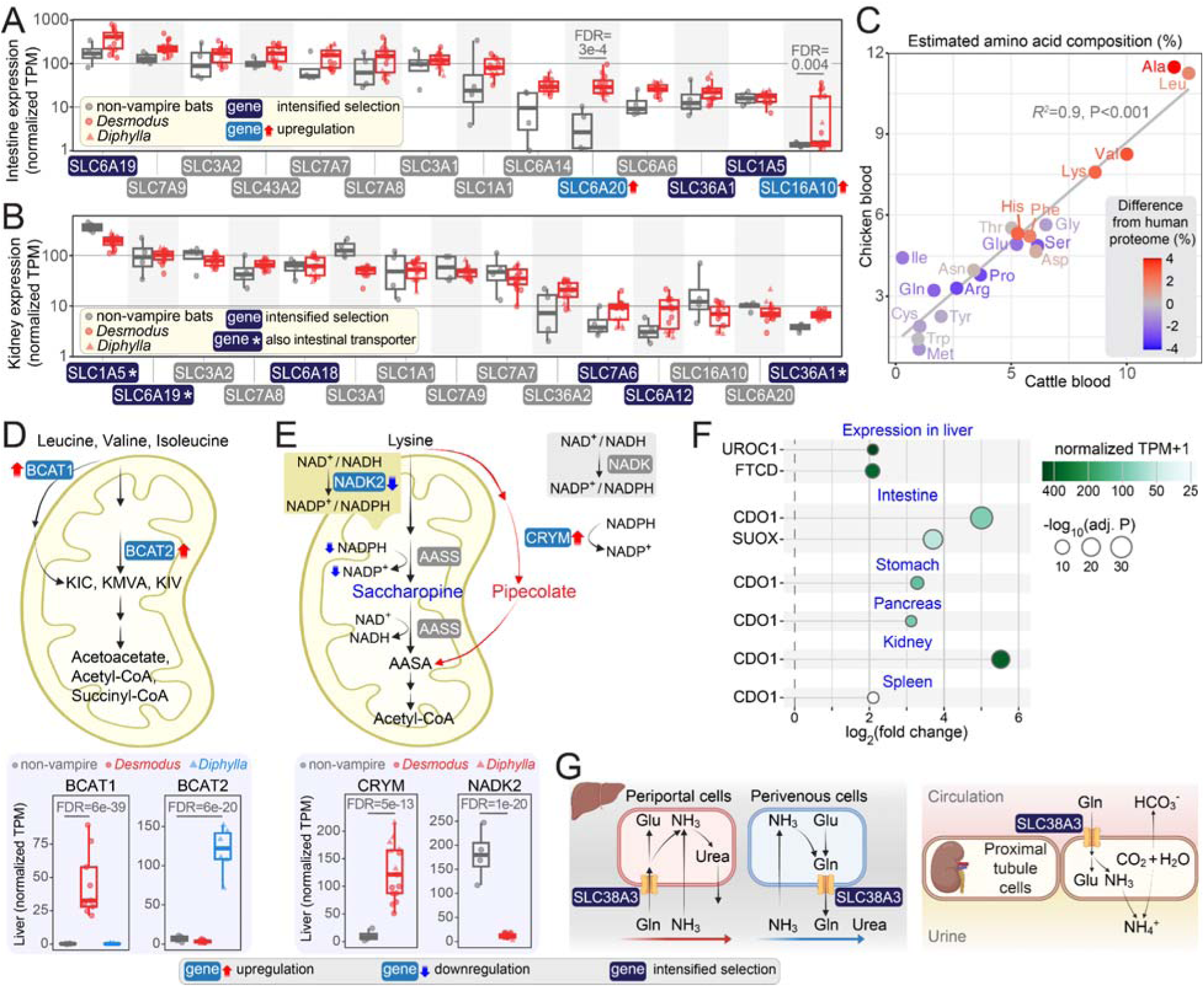
Amino acid transport and catabolism. (A-B) Expression of intestinal (A) and renal (B) amino acid transporters in vampire bats and non-vampire bats (# in Fig. 1A). Genes are ordered by vampire bat expression levels, and genes under intensified selection are highlighted. Asterisks in (B) highlight intensified transporters mediating both intestinal absorption and renal reabsorption. (C) Amino acid frequencies in cattle (X-axis) and chicken (Y-axis) blood, estimated from the combined hemoglobin and albumin protein fractions and weighted by their relative abundance in each species. Color indicates the difference in amino acid frequency relative to the human proteome. (D) Illustration of the initial steps of branched-chain amino acid catabolism, showing the liver upregulation of *BCAT1* in *Desmodus* and *BCAT2* in *Diphylla* at the bottom. (E) Upregulation of *CRYM* (bottom), encoding the key enzyme in the pipecolate lysine-catabolizing pathway. The saccharopine lysine-catabolizing pathway likely has reduced activity, as it depends on mitochondrial NADPH and the mitochondrial NADP-generating enzyme *NADK2* is the fourth most significantly downregulated liver gene (12-fold). (F) Differential expression of genes involved in amino acid catabolism, showing fold changes relative to non-vampire bats (# in Fig. 1A; X-axis), expression levels (normalized TPM, color gradient) and significance (circle sizes). (G) Intensified selection on the glutamine transporter *SLC38A3*, which is highly expressed in the liver and kidney of all studied bats. In the liver, SLC38A3 imports glutamine into periportal hepatocytes for hydrolysis to glutamate and ammonia and subsequent urea-cycle entry, and exports glutamine from perivenous hepatocytes that incorporate scavenged ammonia into glutamine via glutamine synthetase. In the kidney, SLC38A3 imports glutamine into proximal tubule cells, where it is metabolized to generate ammonia for urinary ammonium excretion and bicarbonate, which is returned to the blood for pH restoration.

To prevent amino acid loss in urine, the kidney reabsorbs amino acids using transporters often shared with the intestine (*126*). The three intestinal transporters under intensified selection (*SLC6A19*, *SLC1A5*, *SLC36A1*) also function in renal reabsorption, with *SLC6A19* and *SLC1A5* being the most highly expressed kidney transporters in bats (Fig. 4B). Furthermore, we observed intensified selection on three additional renal transporters: *SLC6A18* (transporting glycine and alanine), *SLC7A6* (arginine and lysine) and *SLC6A12* (alanine) (*126*, *127*). Together, these intensified selection signatures and expression shifts indicate increased importance of intestinal absorption and renal reabsorption of amino acids in vampire bats.

### Upregulated amino acid catabolism

Amino acids are the main energy source for vampire bats (*18*). Our analyses provide molecular insights into this shift in energy metabolism by revealing upregulation and intensified selection on key genes involved in multiple amino acid catabolic pathways, including those metabolizing amino acids abundant in blood (Leu, Val, Lys; Fig. 4C).

Unlike other amino acids, the initial step of branched-chain amino acid (leucine, valine and isoleucine) catabolism does not occur in liver, because this organ normally lacks the transaminases *BCAT1* and *BCAT2* that catalyze the first reaction (*128*, *129*). In contrast to other mammals, we detected liver expression of *BCAT1* in *Desmodus* and *BCAT2* in *Diphylla* (Fig. 4D), suggesting active utilization of branched-chain amino acids as fuel molecules.

Lysine, the fourth most abundant amino acid in blood (*130*), is catabolized via two distinct pathways. The mitochondrial saccharopine pathway predominates in extracerebral tissues, while the pipecolate pathway is restricted to the adult brain (*131*, *132*). However, the saccharopine pathway enzyme AASS is strictly dependent on NADPH (*131*, *132*) and mitochondria of vampire bat liver cells are likely deficient in NADP/NADPH, as the mitochondrial NADP-generating enzyme *NADK2* is severely downregulated (Fig. 4E). Because mutations in human *NADK2* impair lysine degradation and cause hyperlysinemia (*133*), the saccharopine pathway may have reduced activity in vampire bats. Compensating for this, we discovered that *CRYM*, encoding a key enzyme in the pipecolate pathway, is upregulated in both liver and intestine (Fig. 4E). Although CRYM is also NADPH-dependent, it is located in the cytoplasm, where NADPH is likely not restricted in vampire bats (below). Therefore, the pipecolate pathway may have replaced the saccharopine pathway as the major mechanism for lysine degradation in vampire bats.

While histidine is the second rarest amino acid in human whole body proteins (*134*), it is an abundant amino acid in blood (Fig. 4C). Consistent with vampire bats having a greater reliance on histidine catabolism, *UROC1* and *FTCD*, two enzymes in this pathway, are upregulated in the liver (Fig. 4F). Cysteine comprises ∼6% of the residues in mammalian and bird albumin, an abundant blood plasma protein. The rate-limiting cysteine catabolizing enzyme is *CDO1*, which prevents oxidative stress and cytotoxicity caused by abnormally elevated cysteine levels (*135*, *136*). While *CDO1* is highly expressed in the liver of all analyzed bat species, we found an upregulation of *CDO1* in vampire bats in all five non-hepatic tissues examined (intestine, stomach, pancreas, kidney, spleen) (Fig. 4F). Cysteine catabolism produces toxic sulfite. Sulfite oxidase (encoded by *SUOX*) detoxifies sulfite to sulfate (*137*) and is upregulated in the vampire bat intestine. Together, these expression changes suggest an increased role for cysteine catabolism in non-hepatic tissues. Finally, tryptophan is primarily catabolized via the kynurenine pathway, and TDO2 (tryptophan 2,3-dioxygenase), which encodes the first and rate-limiting enzyme of this pathway, is under intensified selection (Supplementary Table 4).

High reliance on amino acid catabolism imposes two systemic challenges: (i) ammonia toxicity and (ii) metabolic acidosis from catabolized ketonic and sulfur-containing amino acids (*138*, *139*). We found that *SLC38A3*, a glutamine transporter essential for both processes (*140–142*), evolved under intensified selection in vampire bats (Fig. 4G). Since the ancestor of phyllostomid bats lost the related *SLC38A5* transporter gene (Supplementary Figure 19), *SLC38A3* may be particularly important for vampire bats to avoid ammonia toxicity and metabolic acidosis.

### Impaired fat digestion

Vampire bats consume an extremely fat-poor diet. We uncovered losses, relaxation, and downregulation of key genes required for fat digestion (Fig. 5A-D, Supplementary Figures 20-21). The *LIPF* gene, encoding the gastric lipase secreted by gastric chief cells, is lost in all phyllostomid bats analyzed in this study. Carboxyl ester lipase (*CEL*) is downregulated in the vampire bat pancreas. Most strikingly, pancreatic triglyceride lipase (*PNLIP*), the primary enzyme responsible for dietary fat digestion, is lost in *Diaemus* and not expressed in the pancreas of *Diphylla* and *Desmodus*. *PNLIP* also exhibits a polymorphic stop codon in *Desmodus*, potentially indicating an early stage of gene loss. Overall, loss or lack of expression of this major dietary fat-digestion enzyme evolved independently in all three vampire bats.

**Figure 5:**
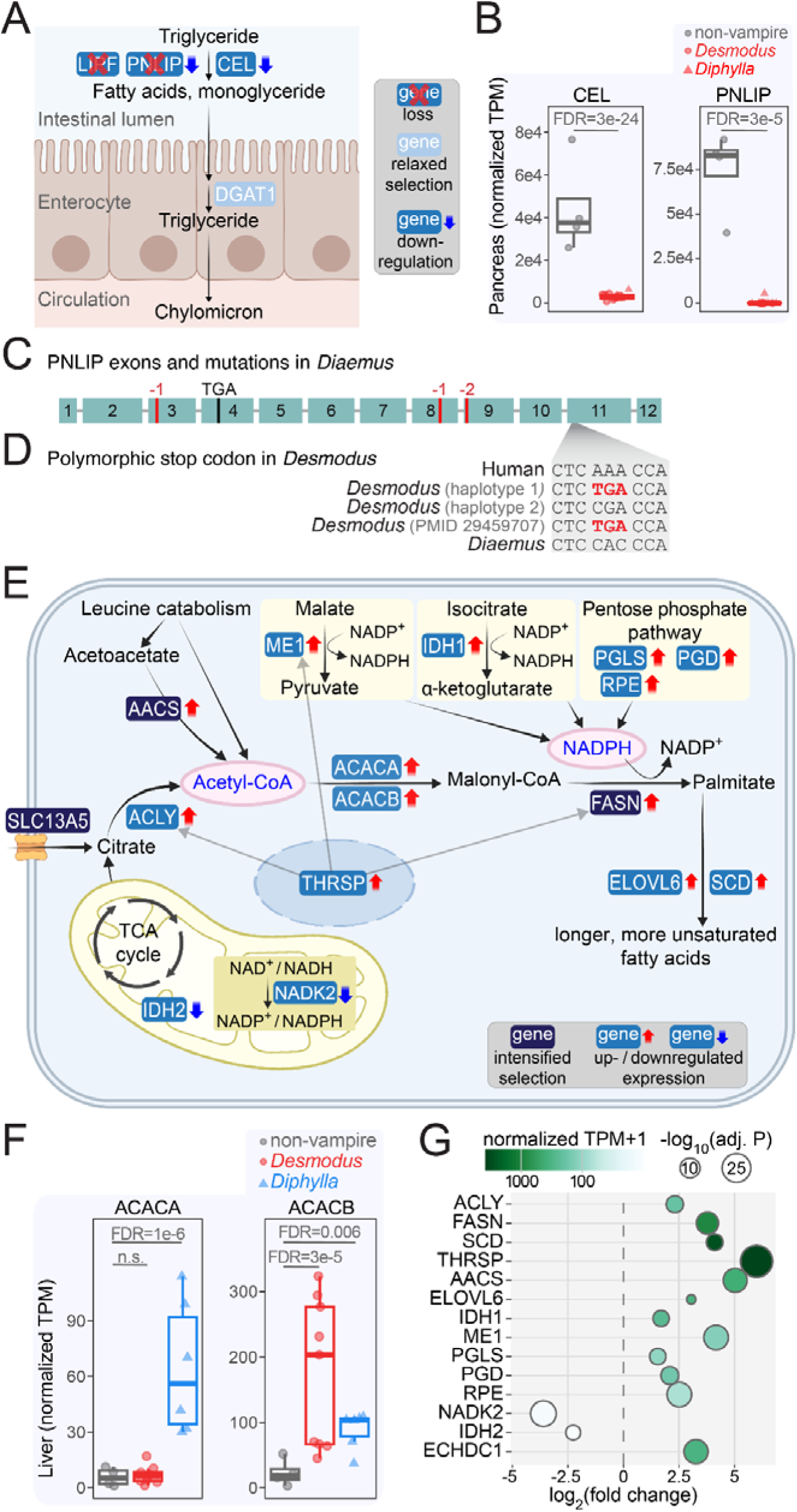
Impaired fat digestion and upregulated fatty acid synthesis. (A) Illustration of intestinal triglyceride digestion and changes in key enzymes in vampire bats. (B) Downregulated pancreas expression of *CEL* and *PNLIP*. (C) Visualization of the coding exons of *PNLIP* in *Diaemus* showing gene-inactivating mutations inferred by TOGA. (D) Polymorphic stop codon in *PNLIP* exon 11, found in haplotype 1 of our *Desmodus* assembly and the previous genome assembly of a different individual (*25*). (E) Illustration of hepatic fatty acid synthesis, highlighting genes with expression changes and intensified selection. (F) Liver expression of *ACACB*, upregulated in both vampire bats, and *ACACA*, upregulated in *Diphylla*. (G) Differential expression of genes involved in fatty acid synthesis in the liver, showing fold changes relative to non-vampire bats (# in Fig. 1A; X-axis), expression levels (normalized TPM, color gradient) and significance (circle size).

Triglyceride digestion results in fatty acids and monoacylglycerol, which enterocytes absorb and reassemble into triglycerides. *DGAT1*, encoding the enzyme catalyzing the final triglyceride reassembly step (*143*), is under relaxed selection in vampire bats (Fig. 5A). Taken together, these results indicate a substantially reduced capacity for fat digestion in adult vampire bats.

Notably, major enzymes responsible for digesting milk fat in infants, pancreatic lipase related protein 2 (*PNLIPRP2*) and *CEL* (*144*, *145*), remain intact in vampire bats. Genes in the bile acid synthesis pathway are also intact and expressed at normal levels (Supplementary Figure 22), which likely ensures the absorption of fat-soluble vitamins (A, D, E, K), cholesterol, and essential fatty acids, and excretion of the heme degradation product bilirubin (*146*, *147*).

### Highly active hepatic *de novo* fatty acid synthesis

Since their diet is extremely poor in fat, vampire bats must synthesize fatty acids from non-lipid precursors. Here we show that *de novo* fatty acid synthesis is drastically upregulated. This process occurs in the cytosol and requires two main substrates, acetyl-CoA and NADPH.

The first substrate (cytosolic acetyl-CoA) is mainly derived from citrate exported from mitochondria. In vampire bats, the enzyme-encoding genes that convert citrate to the fatty acid palmitate (ATP citrate lyase (*ACLY*), acetyl-CoA carboxylase (*ACACB*), fatty acid synthase (*FASN*)) are strongly upregulated in the liver (Fig. 5E-G). *FASN* also evolved under intensified selection and another acetyl-CoA carboxylase, *ACACA,* is upregulated in *Diphylla* (Fig. 5F). *SCD*, which desaturates FASN-generated palmitate, is also upregulated. Consistent with enhanced fatty acid synthesis, *THRSP*, a nuclear factor that induces *ACLY*, *FASN* and malic enzyme (*ME1*, discussed below) expression (*148*, *149*), is the third-most upregulated gene in the vampire bat liver. Additionally, *SLC13A5*, a high-capacity transporter importing extracellular citrate into the cytoplasm (*150*), evolved under intensified selection.

Acetyl-CoA can also be generated from catabolism of blood-abundant leucine (Fig. 4C), which additionally yields the ketone body acetoacetate. In the cytosol, acetoacetate can be converted by AACS into acetoacetyl-CoA (consisting of two acetyl-CoA molecules) to fuel *de novo* fatty acid and cholesterol synthesis (*151*, *152*). We found that *AACS* evolved under intensified selection and is one of the most upregulated genes in the vampire bat liver (36-fold; rank 5, Fig. 5E,G), while being barely expressed in other bats. AACS also contributes to the elongation of polyunsaturated fatty acids via elongases (*153*), and we found the elongase gene *ELOVL*6 upregulated in the vampire bat liver. While the *AACS* pathway plays a minor or condition-specific role in human and mouse (*152*), intensified selection and upregulated *AACS* expression indicate that leucine-derived acetoacetyl-CoA may be an important substrate for fatty acid and cholesterol synthesis in vampire bats.

The second substrate for fatty acid synthesis (cytosolic NADPH) can be generated from three main sources: (i) the pentose phosphate pathway, which generates two NADPH molecules, (ii) IDH1, which converts isocitrate to α-ketoglutarate, and (iii) ME1, which converts malate to pyruvate (*154*). The vampire bat liver exhibits a significant upregulation of *PGLS*, *PGD* and *RPE* in the pentose phosphate pathway, as well as *IDH1* and *ME1*, suggesting that all three sources contribute to increased NADPH production (Fig. 5E,G). Whereas the substrates for the pentose phosphate pathway and *IDH1* are primarily derived from glucose, which is limited in the vampire bat diet, the ME1 substrate (malate) is readily produced by catabolism of abundant dietary amino acids via the urea cycle. This potentially explains why the malate-consuming *ME1* is the most upregulated of these three pathways (17-fold, rank 4). Together, intensified selection signatures and coordinated transcriptional upregulation indicate that fatty acid synthesis is considerably elevated in vampire bats, likely reflecting an adaptation to their fat-poor diet.

The only mechanism of *de novo* NADP(H) generation is the phosphorylation of NAD+, which is executed by *NADK* in the cytosol and by *NADK2* in mitochondria (*155*). While *NADK* is broadly expressed and shows no expression difference in vampire bats, *NADK2*, which is highly expressed in the liver of humans and non-vampire bats, is drastically downregulated in the vampire bat liver (12-fold, rank 4) (Fig. 4E,5E,5G). Furthermore, *IDH2*, encoding the mitochondrial NADP+-dependent enzyme that generates NADPH during isocitrate to α-ketoglutarate conversion, is also downregulated in the vampire bat liver. This suggests a highly unusual situation of low mitochondrial but high cytosolic hepatic NADP(H) levels. Low mitochondrial NADP(H) provides a potential mechanism to inhibit DECR1, the rate-limiting NADPH-dependent enzyme required for β-oxidation of polyunsaturated fatty acids (*156*, *157*), which may indirectly increase the cytosolic NADPH pool and thereby enhance fatty acid synthesis.

The promiscuous FASN enzyme can utilize side products of branched-chain amino acid catabolism, such as methylmalonyl-CoA or ethylmalonyl-CoA, in place of acetyl-CoA, thereby producing branched-chain fatty acids that can be incorporated into phospholipids and perturb membrane fluidity. *ECHDC1*, encoding the enzyme limiting branched-chain fatty acid formation (*158*), is highly upregulated in the vampire bat liver (rank: 12, Fig. 5G), likely because of increased branched-chain amino acid catabolism.

### Decreased reliance on sugar as an energy source

While sugars are a major energy source for many mammals, blood contains extremely low levels of carbohydrates (1%) (*14*, *15*), indicating that vampire bats depend less on sugar as an energy source. Our integrative analysis suggests potential genetic mechanisms underlying vampire bats’ decreased capacity to digest, metabolize, and store sugars.

*SI*, encoding the intestinal brush border sucrase-isomaltase enzyme that cleaves dietary disaccharides into glucose and fructose (*159*), is lost in *Diphylla* and exhibits polymorphic gene-inactivating mutations in *Diaemus* (Fig. 6A, Supplementary Figure 23). The rate-limiting step of aerobic glucose oxidation is catalyzed by the pyruvate dehydrogenase complex (PDC), which converts pyruvate to acetyl-CoA and connects cytosolic glycolysis with the mitochondrial TCA cycle and oxidative phosphorylation (*160–162*). We found *PDHA1* and *PDHB*, which encode the catalytic PDC subunits, to evolve under relaxed selection in vampire bats (Fig. 6B). Notably, mutations affecting the PDC, particularly *PDHA1*, cause lactic acidosis in humans, which can be treated with a ketogenic diet to avoid the glycolytic overproduction of the PDC substrate pyruvate (*160*, *161*, *163*); this dietary intervention resembles the low-sugar vampire bat diet. Genes involved in fructose catabolism (*KHK* and *TKFC*) are also downregulated in the vampire bat liver and kidney (Fig. 6C,D), consistent with the extremely low fructose content of blood, with concentrations even lower than glucose (*164*).

**Figure 6:**
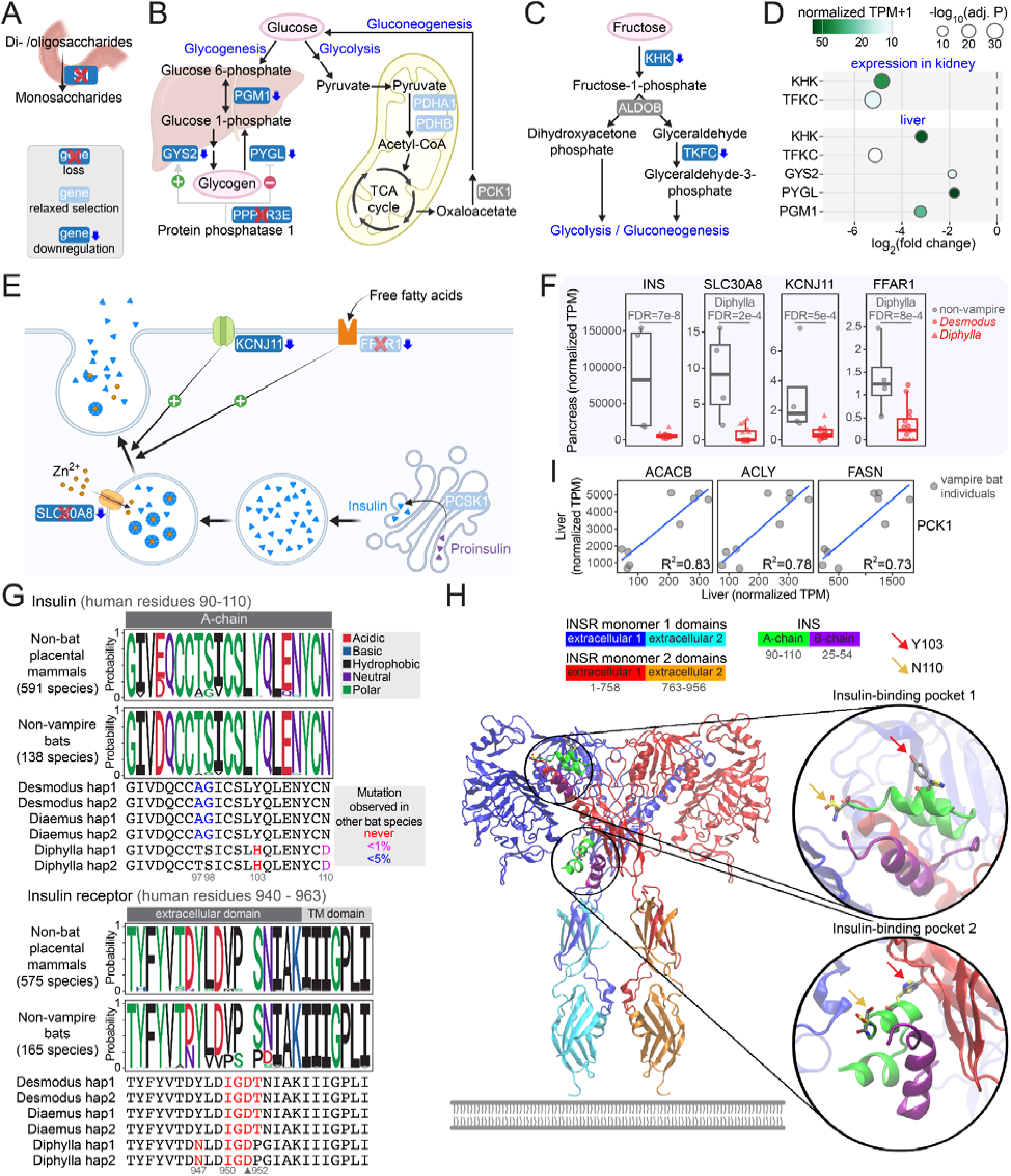
Impaired sugar and insulin metabolism. (A-C) Losses, relaxation and downregulation of genes involved in intestinal sucrose digestion (A), glucose oxidation and glycogen metabolism (B), and fructose catabolism (C). (D) Differential expression of genes involved in sugar metabolism, showing fold changes relative to non-vampire bats (# in Fig. 1A; X-axis), expression levels (normalized TPM, color gradient) and significance (circle size). (E) Sequence and expression changes in genes important for insulin storage and secretion. Schematics as in (A). (F) Expression level of insulin and genes involved in insulin secretion. (G) Mutations in insulin and insulin receptor. Sequence logos represent the relative frequency of residues. The *Desmodus* T97A and S98G occur in <5% of other bat species. The *Diphylla* Y103H and N110D INS mutations occur in no and only one other bat species, respectively. Highlighted mutations in vampire bat *INSR* are never observed in other bats. (H) Cryo-EM structure of the homodimeric insulin receptor extracellular domain bound to insulin (PDB 6PXV). Close-up views highlight insulin-binding pockets, showing the residues differing between human (Y103 and N110) and *Diphylla* (H103 and D110) insulin with carbon atoms colored grey and yellow, respectively. The transmembrane and intracellular domains are unresolved in the cryo-EM structure; a simplified membrane is therefore shown at the base of the extracellular domain.

Glucose is primarily stored as glycogen. Protein phosphatase 1 promotes glycogen synthesis and inhibits glycogen breakdown by dephosphorylating glycogen synthase and glycogen phosphorylase, respectively (*165*, *166*). The regulatory subunit of protein phosphatase 1 encoded by *PPP1R3E*, previously found to be lost in *Desmodus* (*31*), is independently lost in all vampire bats (Supplementary Figure 24). Furthermore, we observed hepatic downregulation of *GYS2*, encoding the liver glycogen synthase, *PYGL*, encoding the liver form of glycogen phosphorylase, and *PGM1* (phosphoglucomutase 1), which interconverts glucose-1-phosphate and glucose-6-phosphate for glycogen synthesis and glycogen breakdown (Fig. 6B,D). Together, these changes provide a plausible molecular explanation for the markedly low glycogen stores in *Desmodus*, which are ∼85% lower than in fruit-eating bats and ∼60% lower than in other mammals fed high-protein diets (*17*, *167*).

### Decreased importance of insulin

Vampire bats have smaller pancreatic islets with a reduced beta cell mass, lower insulin levels than other mammals, and exhibit a reduced glucose-stimulated insulin secretion response (*16*, *168*). Here, we uncover potential molecular underpinnings of their insulin secretion deficiency, revealing that several genes critical for insulin production and secretion are lost, relaxed or downregulated (Fig. 6E-F, Supplementary Figures 25-26). *PCSK1*, encoding the prohormone convertase that processes proinsulin into mature insulin (*169*, *170*), is under strong relaxed selection in vampire bats. *KCNJ11*, encoding a glucose sensitive potassium channel that triggers membrane depolarization and insulin release (*171*, *172*), is downregulated in the vampire bat pancreas. *SLC30A8*, encoding the major beta-cell zinc transporter required for the formation of zinc-insulin crystals (the main insulin storage form) (*173*, *174*), was previously found to be lost in *Desmodus* (*31*). Our analysis revealed an independent loss in *Diaemus* and a downregulation in the *Diphylla* pancreas. Similarly, *FFAR1*, encoding a G-protein-coupled receptor that senses circulating free fatty acids and amplifies glucose-stimulated insulin secretion (*175*, *176*), is lost in *Desmodus* and *Diaemus*, unexpressed in *Diphylla*, and exhibits strongly relaxed selection in vampire bats.

Importantly, we found a pronounced decrease in insulin gene expression in the vampire bat pancreas (Fig. 6F). In addition, the alpha chain of vampire bat insulin exhibits amino acid mutations at otherwise highly conserved positions (97, 98, 103, 110; Fig. 6G). Notably, *Diphylla* exhibits a mutation (Y103H) that occurs in no other bat species. Mutations affecting INS residues 97, 98, and 103 have been reported in human patients with permanent neonatal diabetes (*177*, *178*). The crystal structure shows that insulin Y103 is located at the interaction surface with the receptor (insulin-binding pocket 2; Fig. 6H), suggesting that it could influence insulin conformation and receptor interaction. Interestingly, *INSR*, encoding the insulin receptor, showed a signature of positive selection, driven by several consecutive vampire bat-specific amino acid changes, including an insertion (Fig. 6G). The affected residues lie directly upstream of the transmembrane domain in the extracellular region of INSR, a region critical for insulin-induced structural rearrangements, receptor internalization, and ligand degradation (*179*, *180*). Notably, distinct INSR mutations have also been identified in Mexican tetra cavefish populations, another system exhibiting naturally evolved insulin resistance (*181*). Together, reduced insulin expression and sequence changes in both insulin and its receptor point to attenuated insulin signaling in vampire bats.

Finally, insulin normally suppresses gluconeogenesis while promoting lipogenesis in the liver, leading to an inverse regulation of the two processes. Consistent with insulin deficiency in vampire bats, we observed a positive correlation between the liver expression of *PCK1*, the first-step enzyme of gluconeogenesis, and key lipogenesis genes *ACACB*, *ACLY* and *FASN* (Fig. 6I), suggesting coordinated expression of gluconeogenesis and lipogenesis genes. This resembles the situation in type 2 diabetic patients, where insulin resistance impairs the suppression of gluconeogenesis, leading to both gluconeogenesis and lipogenesis occurring at the same time (*182*). For vampire bats, this regulatory paradigm shift may be beneficial, because unlike most mammals that use gluconeogenesis for energy supply during fasting, they use gluconeogenesis to convert amino acids to glucose after feeding.

### Upregulated heme iron absorption

Compared to other diets, blood contains considerably more iron (about 800-fold higher than a typical human diet (*183*)), primarily as heme-bound iron. Although iron absorption normally occurs in the small intestine, our results suggest that the vampire bat stomach has also acquired this capacity. Specifically, their stomach shows upregulation of the heme transporter genes *SLCO2B1* and *SLC46A1* (*184*, *185*), and heme oxygenase 1 (*HMOX1*), the rate-limiting enzyme that degrades heme to ferrous (Fe^2+^) iron and biliverdin (*186*) (Fig. 7A,B). Notably, *SLCO2B1* and *HMOX1* are also upregulated in the intestine. We also observed increased stomach expression of ferroportin (*SLC40A1*), the only mammalian iron exporter transporting iron from cells into the bloodstream (*187*), and the liver downregulation of hepcidin (*HAMP*), a hormone that lowers circulating iron levels by promoting ferroportin degradation (*188*). Both changes may contribute to the elevated blood iron levels observed in vampire bats (*31*). Finally, increased stomach expression of two ferroxidase enzymes hephaestin (*HEPH*) and ceruloplasmin (*CP*), which convert ferrous (Fe^2+^) to ferric (Fe^3+^) iron, is consistent with increased capacity for transferrin-mediated iron transport in the bloodstream (*189*). In contrast to the upregulation of heme iron-absorbing genes, *CYBRD1*, the plasma membrane ferrireductase required for absorbing free dietary iron (*190*), is ∼53-fold downregulated in the intestine (Fig. 7A,B). Whereas most mammals rely on free iron uptake as the main iron source and thus require *CYBRD1*, the abundance of heme iron likely reduces its importance in vampire bats. Together, these findings suggest that both the stomach and intestine contribute to the absorption and metabolism of dietary heme-bound iron in vampire bats.

**Figure 7:**
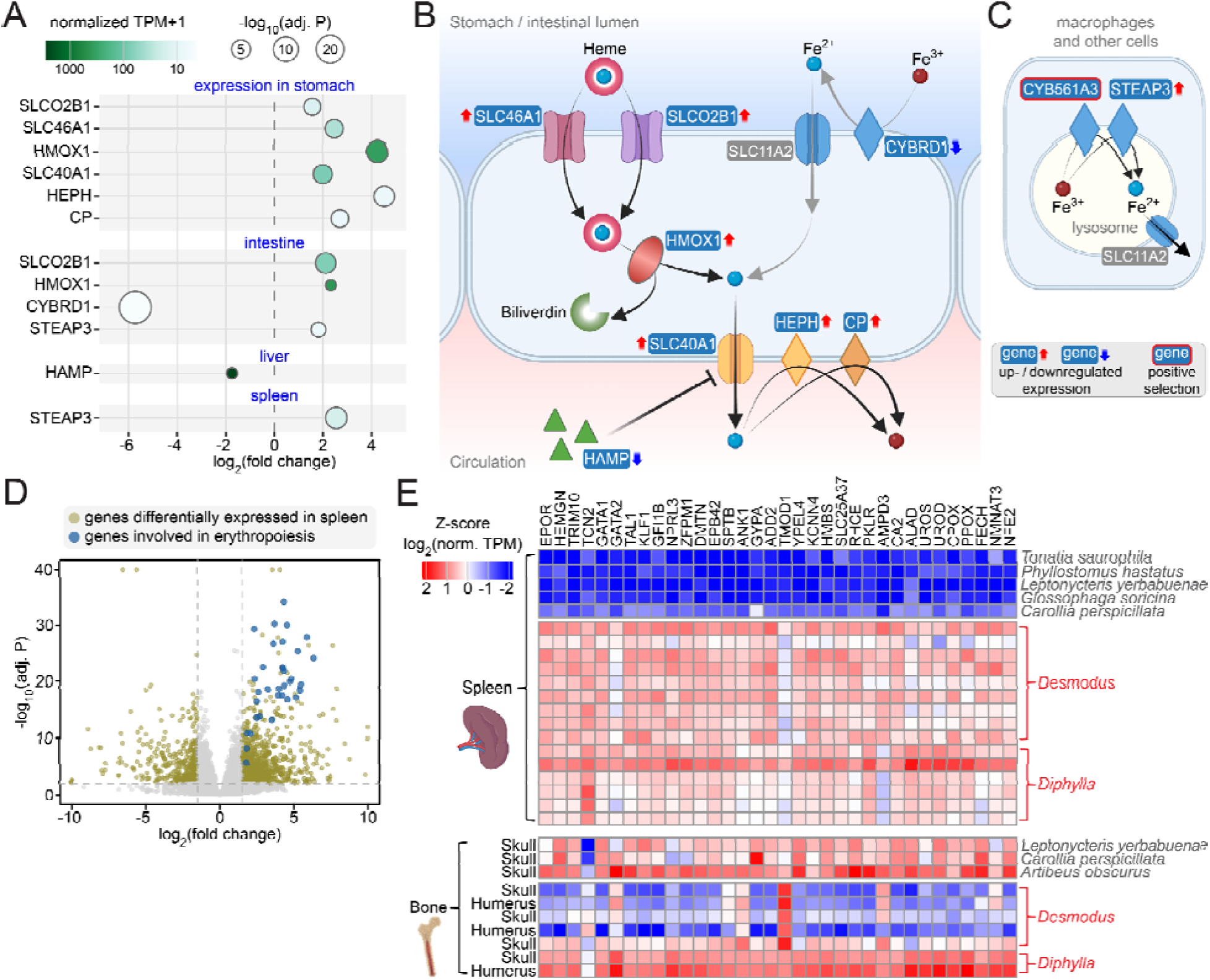
Upregulated heme iron absorption and splenic erythropoiesis. (A) Differential expression of genes involved in iron absorption, showing fold changes relative to non-vampire bats (# in Fig. 1A; X-axis), expression levels (normalized TPM, color gradient) and significance (circle size). (B-C) Illustration of sequence and expression changes of genes involved in heme and free iron absorption (B) and iron recycling (C). (D) Volcano plot showing genes up- and downregulated in the vampire bat spleen. Genes directly involved in erythropoiesis are highlighted in blue. (E) Expression levels of key erythropoietic genes in adult spleen and bone of vampire (red font) and control bats (black font), visualized as per-gene Z-scores of log_2_(normalized TPM + 1).

Two intracellular ferrireductases involved in iron recycling exhibit notable changes in vampire bats: *CYB561A3* evolved under positive selection on the ancestral vampire bat branch and *STEAP3* is upregulated in the intestine and spleen (Fig. 7A,C). Both enzymes generate bioavailable ferrous iron in macrophages that engulf defective erythrocytes and recycle iron to sustain erythropoiesis (*191*). In addition, *STEAP3* is found in endosomes of erythroid cells, where it is required for efficient erythropoiesis (*192*, *193*).

While a previous study described an expansion of iron-storing ferritin genes in *Desmodus* (*25*), our reanalysis revealed a single ferritin heavy chain ortholog across bats as well as an ancient, non-vampire bat-specific duplication of the ferritin light chain gene (Supplementary Figures 27-29, Supplementary Table 22). We attribute the previously-reported ferritin expansion to processed pseudogene copies. Importantly, ferritin processed pseudogenes occur in all analyzed phyllostomid and outgroup bats, hence are unlikely to be related to the iron rich vampire bat diet.

Overall, our findings are consistent with vampire bats enhancing, rather than limiting, the absorption of dietary heme-bound iron.

### Splenic erythropoiesis in vampire bats

Red blood cell production (erythropoiesis) requires large amounts of iron, vitamin B9, and vitamin B12 (*194*); all appear to be absorbed at elevated levels in vampire bats. In most placental mammals, erythropoiesis occurs primarily in the bone marrow, whereas the spleen supports erythropoiesis only during fetal development or under stress and pathological conditions in adults (*195*, *196*).

Strikingly, we found strong transcriptomic evidence that the spleen is a major erythropoietic organ in adult vampire bats. Genes upregulated in the spleen are highly enriched for heme and hemoglobin biosynthesis, and erythrocyte differentiation (Fig. 1I). Of the 50 most significantly upregulated genes, 25 are directly involved in erythropoiesis or erythrocyte function (Fig. 7D,E). These include genes required for erythroid differentiation (erythropoietin receptor (*EPOR*), hemogen (*HEMGN*), *TRIM10*, *TCN2*, *GATA1*, *GATA2*, *TAL1*, *KLF1*, *GFI1B*, *NPRL3*, *ZFPM1*), erythrocyte membrane stability (*DMTN*, *EPB42*, *SPTB*, *ANK1*, *ADD2*, *TMOD1*, *YPEL4*) and cell volume regulation (*KCNN4*), iron uptake and hemoglobin synthesis (*HMBS* and *SLC25A37*, encoding mitoferrin-1), blood group antigen expression (*RHCE*), and erythrocyte metabolism (*PKLR*, *AMPD3*, *CA2*) (*197–200*). Nearly all heme biosynthesis genes (*ALAD*, *HMBS*, *UROS*, *UROD*, *CPOX*, *PPOX*, *FECH)* are significantly upregulated. Furthermore, *NMNAT3*, encoding the main NAD+ producing enzyme required for erythrocyte glycolysis and energy metabolism (*201*, *202*), is upregulated in the spleen. Transcriptional regulators of megakaryocyte and platelet development (*GFI1B*, *NFE2*) are also upregulated.

To test whether bone marrow remains an active site of erythropoiesis in vampire bats, we generated ten additional RNA-seq datasets from skull and humerus bones, which are typical sites of adult erythropoiesis, in *Desmodus*, *Diphylla*, and three non-vampire bat species (Supplementary Table 5). As expected, non-vampire bats showed high expression of the erythropoietic gene program (Fig. 7E). In vampire bats, expression of erythropoietic genes in bone marrow was generally low in *Desmodus* but high in *Diphylla*, suggesting that the spleen became the primary erythropoietic organ in *Desmodus*, whereas both spleen and bone marrow likely contribute to erythropoiesis in *Diphylla*. Although the reasons for splenic erythropoiesis in vampire bats remain to be investigated, the spleen may provide a favorable site by spatially coupling macrophage-mediated erythrocyte clearance and iron recycling with erythrocyte production within a single organ.

## Discussion

Our high-quality reference genomes of all three vampire bat species enabled integrative comparative genomic and transcriptomic analyses that revealed coordinated molecular changes associated with obligate sanguivory in vampire bats, the only tetrapods to have evolved this extreme diet. We identified molecular mechanisms potentially contributing to a radical transformation of the vampire bat stomach from an acidic, digestive organ into an enlarged, non-acidic chamber that exhibits an altered mucus profile and is specialized for water, electrolyte, vitamin and heme absorption. Upregulation of connective tissue-related genes is consistent with our histological evidence of increased connective tissue content, which may provide structural support for extreme stomach expansion during a single feeding bout.

The extreme dietary specialization of vampire bats is mirrored by extensive molecular remodeling in key metabolic pathways. Consistent with their fat-poor diet, digestive lipases are lost or evolve under relaxed selection, whereas lipogenic enzymes and genes generating the lipogenic substrates (acetyl-CoA and NADPH) evolve under intensified selection and are transcriptionally upregulated. Low dietary sugar content is reflected by loss, relaxation, or downregulation of genes involved in carbohydrate metabolism and insulin signaling, revealing molecular insights into the evolutionary origins of natural insulin deficiency in vampire bats. Notably, the naturally low-fat, low-sugar vampire bat diet parallels dietary interventions used to manage human metabolic disorders caused by deficiencies in *DGAT1*, *PDHA1*, or *INSR* (*163*, *203*, *204*), genes that are also altered in these bats. Adaptation to a protein-rich diet is supported by independent duplications of trypsin genes, as well as by intensified selection and upregulation of genes involved in intestinal absorption, renal reabsorption, and amino acid catabolism. Our results also suggest that key aspects of amino acid catabolism in vampire bats diverge from those of other mammals, including (i) hepatic catabolism of branched-chain amino acids, (ii) expression changes consistent with a shift in lysine catabolism, with the normally brain-restricted pipecolate pathway expanded to liver and intestine, and (iii) a role of leucine-derived acetoacetyl-CoA as a substrate for fatty acid synthesis. Although many molecular changes are shared between *Desmodus* and *Diphylla*, we also observed changes in distinct but functionally equivalent genes, such as *BCAT1*/*BCAT2* and *ACACA*/*ACACB*, suggesting different genetic routes to similar metabolic outcomes.

Most genes lost in vampire bats are associated with gastric mucosal function, gastrointestinal motility, and nutrient digestion. Many of these losses likely reflect relaxed selection on functions that became less important with their exclusive blood diet. In line with the “use it or lose it” principle (*205*), most of these genes were lost independently in vampire bats, and our haplotype-resolved assemblies revealed polymorphic inactivating mutations in several genes (*CTSE*, *MLNR*, *SI*, *PNLIP*), indicative of ongoing gene erosion. However, gene loss can also confer adaptive benefits (*4*, *205–207*). For instance, the loss of the 24S-hydroxycholesterol-degrading enzyme *CYP39A1* may have contributed to their exceptional social behavior and cognitive abilities (*31*). Consistent with this, *CYP39A1* is among the few genes already inactivated in the common ancestor of vampire bats (Supplementary Figure 30).

Comparative genomic analyses uncover gene sequence changes such as gene losses, duplications, and signatures of selection, but they often highlight individual loci and therefore provide a fragmented view of relevant genomic changes. Our study shows that incorporating comparative transcriptomic data across multiple organs adds a complementary layer by revealing coordinated sequence and expression changes across pathways and molecular processes. This integrative approach was essential for obtaining a comprehensive view of the molecular changes associated with sanguivory in vampire bats. More generally, our work demonstrates how genome-wide, integrative analyses can yield molecular insights into the evolution of adaptive traits.

## Materials and Methods

### Sample collection, ethical statements and permits

Tissue samples for genome sequencing of the three vampire bat species were obtained from the Royal Ontario Museum mammal collection, where samples were kept frozen since their collection. Samples of a *Desmodus rotundus* male individual (F63247) were collected at Sabajo, Para, Suriname on 2017-09-26. Samples of a *Diaemus youngii* male individual (F50745) were collected at the Ceiba Biological Center in Guyana on 2001-10-16. Samples of a *Diphylla ecaudata* male individual (F37596) were collected at the Parque Nacional Yasuni (66 km S of Pompeya Sur) in Napo, Ecuador on 1995-10-15. All three vampire bat samples are from male individuals caught in the wild.

Samples of *Carollia perspicillata* for genome sequencing were obtained on 2018-02-21 from a captive breeding colony in Frankfurt, Germany, in accordance with the guidelines of the Darmstadt Regional Council. The colony was established in 1980 from founding animals that were caught at Hacienda La Pacifica, Guanacaste in Costa Rica. Samples of a *Glossophaga soricina* juvenile were obtained on 2018-02-20 from a captive breeding colony in Berlin, Germany, that was established in 1987 from founding animals that were caught in central Mexico. The orphaned juvenile had been permanently abandoned by the mother and therefore had to be euthanized, in compliance with the institutional animal welfare regulations. We observed half-coverage of PacBio reads only for scaffold 9, which aligns to the human X chromosome, indicating that the juvenile was a male. A liver sample of a *Micronycteris megalotis* male adult individual was collected from the wild on 2019-03-14 in Santa Maria, Boyaca, Colombia.

All samples were flash frozen in liquid nitrogen after being collected and stored at -80℃ until further processing. The ethical statements of collecting and processing tissue samples for each species followed the procedures required by the following permits:

- *Diphylla ecaudata* (ROMM105272) – Permit number 048IC from the Instituto Ecuatoriano Forestal y de Áreas Naturales y Vida Silvestre
- *Diaemus youngii* (ROMM113765) – Permit number 20011025 from the Guyana Environmental Protection Agency, Natural Resources Management Division
- *Desmodus rotundus* (ROMM126221) – Permit number 12049 from the Suriname Forest Service, Nature Conservation Division
- *Micronycteris megalotis* (Mm_FF)*–* Permit number DECRETO 1076 DE 2015 (Mayo 26) issued by the The Ministry of Environment and Sustainable Development of Colombia (Spanish: Ministerio de Ambiente y Desarrollo Sostenible de Colombia)

To perform transcriptomics, we obtained wild *Desmodus* and *Diphylla* individuals from a farm in Divinesia, Minas Gerais, Brazil, and a wild *Artibeus obscurus* individual from Vicosa, Minas Gerais, Brazil under the sampling and capturing permit SISBIO 91340-3. The access of these genetic resources from Brazil is registered at SISGEN under the number A3D3AA5. A *Leptonycteris yerbabuenae* individual was obtained from a captive breeding colony in Tübingen, Germany that was established in 1988 from founding animals that were caught in Mexico.

To perform histology, stomach pH measurements, and gastrointestinal motility assessments, we obtained wild *Desmodus* and *Diphylla* individuals from a farm in Divinesia, Minas Gerais, Brazil, wild *Artibeus lituratus* and *Myotis nigricans* individuals from Vicosa, Minas Gerais, Brazil (permit number CEUA 33/2025). Captive *Phyllostomus discolor* individuals were obtained from a captive breeding colony at St. Andrews, UK.

### Transcriptome sequencing

All tissue samples for transcriptome sequencing were immediately stored in RNAlater to ensure RNA preservation. The only exceptions are the *Leptonycteris yerbabuenae* skull and spleen samples, which were flash-frozen. Total RNA was extracted using the RNeasy Plus Mini Kit (Qiagen) according to the manufacturer’s instructions. RNA concentration was measured using fluorometric quantification, and RNA integrity was assessed prior to library preparation using an Agilent TapeStation instrument. Polyadenylated RNA was enriched from total RNA by oligo-dT priming and reverse transcribed into first- and second-strand cDNA followed by indexing adaptor ligation. Strand-specific RNA sequencing libraries were prepared with the Illumina Stranded mRNA Library Prep protocol (Illumina), according to the manufacturer’s instructions. Library quality and fragment size distribution were evaluated, libraries were quantified, and equimolar amounts of each library were pooled for sequencing. Libraries were sequenced on an Illumina platform using paired-end sequencing (2x150 bp), generating approximately 20 million read pairs per sample.

### Genome sequencing

For *Diaemus* and *Diphylla*, we generated PacBio HiFi and Dovetail Omni-C data according to the following protocols. To extract high molecular weight genomic DNA, we followed the protocol of (*208*). DNA concentration and DNA fragment length were assessed using the Qubit dsDNA BR Assay kit on the Qubit Fluorometer (Thermo Fisher Scientific) and the Genomic DNA Screen Tape on the Agilent 2200 TapeStation system (Agilent Technologies), respectively. For each species, one SMRTbell library was prepared according to the instructions of the SMRTbell Express Prep Kit v2.0 with a final size selection using AMPure PB beads. Quality control of the final libraries was performed on the Agilent 2200 TapeStation system (Agilent Technologies). Total input DNA was approximately 1.5 µg per library. The library of each species was loaded three times on the Sequel System IIe in CCS mode, using 30 hours of movie time with two hours of pre-extension and sequencing chemistry v2.0. The on-plate concentration for each Sequel IIe run was 80 pM using diffusion loading. For *Desmodus*, we used existing HiFi and Omni-C data (*31*), and to further improve the assembly, we generated additional HiFi data from one SMRT cell, constructing the SMRTbell library according to the instructions of the SMRTbell Express Prep Kit v3.0.

To generate long-range data for *Diaemus* and *Diphylla* for scaffolding, we used the Dovetail Omni-C kit (Dovetail Genomics, Scotts Valley, CA, USA) following the manufacturer’s protocol (manual version 1.2 for mammalian samples). We used approximately 30 mg (*Diphylla*) and 60 mg (*Diaemus*) of liver tissue from the same individual used for PacBio sequencing. Fragment size distribution and concentration of the Dovetail Omni-C libraries were assessed using the TapeStation 2200 (Agilent Technologies) and the Qubit Fluorometer and Qubit dsDNA HS reagents Assay kit (Thermo Fisher Scientific, Waltham, MA), respectively. The libraries were sequenced on the NovaSeq 6000 platform at Novogene (UK) using 150 bp paired-end sequencing to about 60X coverage totaling 120 Gb.

For *Micronycteris megalotis*, HMW gDNA was extracted and PacBio HiFi data was generated as follows. HMW gDNA was extracted from liver tissue according to the Circulomics Nanobind Tissue Big DNA Kit (PacBio / Circulomics, Handbook v1.0 (11/2019)). Briefly, snap-frozen liver tissue was finely minced with a scalpel and homogenised on ice in chaotropic buffer using the Tissue Ruptor (Qiagen). Tissue lysis was performed by adding Proteinase K and cell debris was removed by centrifugation. The released gDNA was bound to a Nanobind disk after the addition of salt buffer and isopropanol. After several washes, the gDNA was eluted from the Nanobind disk. Genomic DNA concentration was determined using the Qubit dsDNA BR (Broad Range) assay kit (ThermoFisher, USA). The purity of the sample was measured using the NanoDrop by evaluating the curve shape and the 260/280 nm and 260/230 nm values. Pulse-field gel electrophoresis (PFGE) using the Pippin Pulse^TM^ instrument (SAGE Science) revealed HMW DNA molecules <200 kb in length. The HMW gDNA was post-purified with Ampure beads and entered into library preparation using the PacBio HiFi library preparation protocol “Preparing HiFi Libraries from Low DNA Input Using SMRTbell Express Template Prep Kit 2.0”. Briefly, gDNA was sheared to 20 kb using the MegaRuptor device (Diagenode) and 3.9 µg of sheared gDNA was used for library preparation. Fragments shorter than 3 kb were removed using Ampure beads size selection. The size-selected library was prepared for loading according to the instructions generated by the SMRT Link software and the ‘HiFi Reads’ application. The HiFi library was sequenced on 8M SMRT™ Cells for 30 hours with the Sequel^®^ II Binding 2.0 and the Sequel^®^ II Sequencing 2.0 chemistry on the Sequel^®^ II. Circular Consensus sequences were called using PacBio ccs (v6.0.0).

For *Carollia perspicillata* and *Glossophaga soricina*, we generated PacBio CLR and 10x Genomics data as follows. High molecular weight (HMW) genomic DNA (gDNA) was extracted according to the Bionano Prep Animal Tissue DNA Isolation Soft Tissue Protocol (document number 30077, document revision C). Briefly, 20-40 mg of snap-frozen spleen (*Carollia perspicillata*) and liver (*Glossophaga soricina*) tissues were thawed and chopped into small pieces. Tissues were homogenised on ice in a homogenisation buffer using a tissue grinder to isolate nuclei. These were washed in ice-cold ethanol, enriched by centrifugation and embedded in 2% agarose plugs. The plugs were further treated with Proteinase K and RNase A, and then washed with 1X Wash Buffer from the kit and home-made 1X TE buffer (pH 8.0). Ultra-long gDNA was recovered from the plugs and finally purified by drop dialysis against the TE buffer (pH 8.0). gDNA concentration was determined using the Qubit dsDNA BR (Broad Range) assay kit (ThermoFisher, USA). Sample purity was measured with the NanoDrop by evaluating the shape of the curve and the 260/280 nm and 260/230 nm values. The integrity of the HMW gDNA was determined by pulse-field gel electrophoresis using the Pippin Pulse^TM^ device (SAGE Science). The majority of the long and ultra-long gDNA was between 50 and 500 kb in length. All pipetting steps of the ultra-long gDNA were performed very carefully with wide-bore pipette tips.

To sequence PacBio CLR for both species, long insert libraries were prepared as recommended according to the ‘Guidelines for preparing size-selected >30 kb SMRTbell templates making use of the SMRTbell express Template kit 1.0. Briefly, ultra-long gDNA was sheared to 60 kb fragments with the MegaRuptor1 device (Diagenode) and 4-5 µg of sheared gDNA was used for library preparation. The PacBio SMRTbell library was size-selected for fragments larger than 30 kb (*Carollia perspicillata*) and larger than 25 kb (*Glossophaga soricina*) with the BluePippin^TM^ device according to the manufacturer’s instructions. All size-selected libraries were sequenced for 10 hours on the Pacbio Sequel® on 1M SMRT™ cells v2 using the Sequel® Binding 2.1 and the Sequel® Sequencing 2.1 v2 chemistry at the DcGC, Germany.

Ultra-long gDNA from *Carollia perspicillata* and *Glossophaga soricina* was used for 10x genomic linked read sequencing according to the manufacturer’s instructions (10x Genomics Chromium Reagent Kit v2, revision B). Briefly, 1 ng of ultra-long gDNA was loaded into 10x Genomics GEM droplets (Gel Bead-In-EMulsions) using the Chromium device. Genomic DNA molecules were amplified in these individual GEMS in an isothermal incubation using primers containing a specific 16 bp 10x barcode and the Illumina R1 sequence. After breaking the emulsions, pooled amplified barcoded fragments were purified, enriched and used for Illumina sequencing library preparation, as described in the protocol. Sequencing was performed either on a Nextseq 500 High Output flow cell using the 2x150 cycles paired-end regime plus 8 cycles of i7 index at the DcGC, Germany and on a NovaSeq 6000 S1 flow cell using the same sequencing regime at the MPI of Molecular Genetics, Germany.

Chromatin conformation capture (Hi-C) was performed using the Arima Hi-C kit for *Carollia perspicillata* and *Glossophaga soricina* and Arima Hi-C High Coverage kit for *Micronycteris megalotis* according to the Arima documents (User Guide for Animal Tissues, part number A160132 V00 and part number A160162 v00). Briefly, 75 mg of flash-frozen liver tissue from *Carollia perspicillata*, 40 mg flash-frozen brain tissue from *Glossophaga soricina*, and 15 mg of flash-frozen muscle tissue from *Micronycteris megalotis* were chemically crosslinked. The crosslinked genomic DNA was digested with a cocktail of two (*Carollia perspicillata* and *Glossophaga soricina*) and four (*Micronycteris megalotis*) restriction enzymes. The 5’-overhangs were filled in and labelled with biotin. Spatially proximal digested DNA ends were ligated, and the ligated biotin-containing fragments were enriched and used for Illumina library preparation according to the Arima user guide for library preparation using the Kapa Hyper Prep kit (Arima document part number A160139 v00). The barcoded Hi-C libraries were run on S2 and S4 flow cells of an Illumina NovaSeq 6000 at 2x150 cycles at the DcGC, Germany.

### Genome assembly

For the three vampire bats, we produced haplotype-specific assemblies from HiFi reads using Hifiasm (for *Desmodus* v.0.16.1, *Diaemus* v.0.16.1, *Diphylla* v.0.19.3), integrating Hi-C reads for phasing (*32*). Both haplotypes were subject to filtering procedure using Foreign Contamination Screen (FCS) (*209*). Contamination filtered assemblies were then polished to remove unambiguous heterozygous sites. Specifically, we mapped HiFi reads back to each assembly with minimap2 v.2.26 (*210*), removed duplicates using picard MarkDuplicates tool v.3.1.0 (http://broadinstitute.github.io/picard), and called variants using DeepVariant v1.5.0 (*211*). We then filtered for sites with genotype 1/1 and a ‘PASS’ filter value, meaning that all or nearly all reads support an alternative sequence at this position and passed DeepVariants internal filters. Finally, we corrected corresponding nucleotide sites in the assembly with bcftools consensus v.1.13 (*212*).

To scaffold the polished haplotype assemblies of vampire bats, we first mapped the Omni-C reads to each of the two assemblies independently using chromap v.0.2.5 (*213*) and then used YaHS v1.1a (*33*). Finally, we performed manual curation of the two scaffolded haplotypes jointly analyzing them in PretextView v.0.2.5 (https://github.com/wtsi-hpag/PretextView). To this end, we remapped the Hi-C reads to both haplotype assemblies concatenated together using again chromap but allowing for multi-mapping reads (-q 0) to avoid discarding information in regions identical between two haplotypes. We also identified telomeric sequences with tidk v.0.2.31 (https://github.com/tolkit/telomeric-identifier) and where necessary corrected wrong contigs orientations to have telomeres in the ends of resulting scaffolds. To finalize changes made via manual dual haplotype curation, as recently proposed by the Vertebrate Genome Project and Darwin Tree of Life Project, we used the rapid curation framework (https://gitlab.com/wtsi-grit/rapid-curation/-/tree/main) from the Genome Reference Informatics Team (*214*).

For *Micronycteris megalotis*, we generated a primary assembly. Contigs were assembled using PacBio HiFi reads and Hifiasm (v0.15.4-r347) (*32*) with argument -l2. Retained haplotigs were removed from the contig assemblies using Purge_Dups (v1.2.6, parameter -e) (*215*). For scaffolding, Hi-C reads were first mapped to the contig assembly using bwa (v0.7.17, parameter -B8) (*216*) and the Arima mapping pipeline (https://github.com/VGP/vgp-assembly/blob/master/pipeline/salsa/arima_mapping_pipeline.sh). The resulting bam file was used as input for SALSA2 (v2.3, parameters -e GATC,GANTC,CTNAG,TTAA -s 2000000000) (*217*). To manually curate the scaffolded assembly, Hi-C reads were mapped using bwa (v0.7.17, parameter-SP5M), followed by filtering, sorting and deduplication with pairtools (v0.3.0). The resulting pairsam file was then split to bam and converted to .cool and .mcool files using pairx (v0.3.7) and cooler (v0.8.11). Finally, the resulting map was visualised using HiGlass (v1.11.7) and missing or false joins were corrected manually. To close gaps in the scaffolded assembly, we mapped raw subreads to the genome using pbmm2 (v1.7.0, parameters --preset SUBREAD -N 1). Consensus sequences 2 kb upstream and downstream of each gap were then called using PacBio’s gcpp (v2.0.2) and for gap regions which were filled entirely with capital letter substitutions and no gaps (i.e. a high confidence gap fill), the sequences were inserted into the genome around the gap sequences. To correct base errors, the PacBio HiFi reads were mapped using pbmm2 (v1.7.0, parameters: --preset CCS -N 1) and variants were called using DeepVariant (v1.2.0, parameter --model_type=PACBIO) (*211*). The resulting vcf was then filtered with bcftools (v1.16) view with parameter ‘FILTER=“PASS” && GT=“1/1”’ and finally a consensus sequence called with bcftools consensus.

Mitochondrial assemblies were created using MitoHiFi (v2.0) (*218*), but MitoHiFi failed to assemble a mitochondrial genome from the *Diaemus* and from the *Micronycteris* HiFi data. To estimate the assembly base accuracy for the HiFi-based assemblies, we used Merqury (*35*) with the HiFi reads. It should be noted that the QV estimates represent an upper bound as they are based on the HiFi reads used for assembly.

For *Carollia perspicillata* and *Glossophaga soricina*, we generated primary genome assemblies using the DAmar v1 pipeline (https://github.com/MartinPippel/DAmar/blob/master-v1), which orchestrates all assembly steps, including contig building, error correction, and scaffolding. Specifically, the initial contig assembly was performed with the MARVEL assembler (*219*) and the PacBio CLR reads. MARVEL employs a two-phase read-correction procedure that maintains the integrity of long PacBio CLR reads during assembly. Haplotypic duplications were identified and removed with Purge_Dups v1.0.1 (*215*). To increase base level accuracy, we conducted two rounds of error correction using the PacBio data and gcpp (previously known as Arrow; git commit 974589f; https://github.com/PacificBiosciences/gcpp). Subsequently, two additional rounds of error correction were performed using 10x Genomics linked-read data. To this end, 10x linked reads were mapped to the genome using LongRanger v2.2.2 (https://github.com/10XGenomics/longranger) and variants were called using freebayes v1.3.2 (https://github.com/freebayes/freebayes/) with argument “-g 600” to ignore regions with coverage over 600X. The variants were filtered using bcftools view (v1.12) with argument: ‘-i’ QUAL>1 && (GT=“AA” || GT=“Aa”)’ and consensus called using bcftools consensus with argument: ‘-Hla’.

Scaffolding of the *Carollia* and *Glossophaga* assemblies was carried out in two stages. First, Bionano optical map data were used with Bionano Solve (version Solve3.3_10252018; https://bionano.com/software-downloads/). Second, the resulting scaffolds underwent further scaffolding using Hi-C data and SALSA2 (git commit 974589f) (*217*). The assembly was phased using LongRanger v2.2.2 and the 10x linked-read data. To correct scaffolding errors (missing or false joins), we manually curated the assembly by using HiGlass and a HiC map generated with https://github.com/VGP/vgp-assembly/blob/master/pipeline/salsa/arima_mapping_pipeline.sh.

To assess gene completeness, we used compleasm v0.2.7 (https://github.com/huangnengCSU/compleasm) (*44*) with the BUSCO mammalia_ODB12 set of 12,277 near-universally conserved mammalian genes and the TOGA classification of ancestral genes (below).

### Repeat masking

To align newly-sequenced genomes, we generated for each genome a *de novo* repeat library assembly using RepeatModeler 2.0.5 (*220*). We then used the resulting library and RepeatMasker v.4.0.9 (parameters: -engine crossmatch -s) to soft-mask the genome.

### Pairwise genome alignments

To infer orthologous genes with TOGA (below), pairwise alignments between a reference and a query genome are needed. We used the human (hg38) assembly as a reference and generated pairwise genome alignments to the newly-sequenced genomes and other species. Briefly, we used LASTZ 1.04.15 (*221*) with parameters (K = 2400, L = 3000, Y = 9400, H = 2000, default scoring matrix) that are sensitive enough to detect alignments between orthologous exons across placental mammals (*222*). The resulting local LASTZ alignments were then chained using axtChain(*223*) (default parameters, except setting linearGap=loose), followed by using RepeatFiller (*224*) (default parameters) to add missed repeat-overlapping local alignments to the alignment chains and chainCleaner (*225*) (default parameters, except setting minBrokenChainScore = 75,000 and -doPairs) to improve alignment specificity.

### Orthology inference and codon alignments

To identify orthologous genes, we used TOGA v1.1.4 (*39*) (https://github.com/hillerlab/TOGA) with the human hg38 genome and GENCODE V38 annotation (*226*) as the reference and the generated pairwise alignment chains to annotate coding genes and infer orthologs across all considered bat genomes.

Multiple codon alignments were used for phylogenomics and screens for selection signatures. To this end, we extracted 1:1 orthologs from TOGA, taking the longest transcript of each gene. Leveraging that TOGA is aware of exon-level orthology (*39*), we used an exon-by-exon strategy that aligns each orthologous exon individually with MACSE v2 (*227*) and default parameters, followed by concatenating exon alignments into complete codon alignments.

### Phylogenetic and divergence time estimation

To generate a phylogeny that includes both newly sequenced and existing genomes, we used codon alignments of 1:1 orthologous genes. We applied a coalescent-based approach to infer gene trees from each alignment, which were then used to reconstruct a species tree. We included only orthologs classified by TOGA as I (intact), PI (partially intact), or UL (uncertain loss). We excluded codon alignments that were missing sequences for any vampire bat or for more than three non-vampire bat species, and we removed codons that were present in fewer than 50% of species. With these filters, we obtained codon alignments for a total of 15,481 orthologous genes.

For each gene alignment, we inferred maximum-likelihood trees with RAxML v.8.1.16 (*228*) under a GTR+GAMMA model, using three independent replicates and the rapid hill climbing algorithm. Then, we used ASTRAL v.5.5.9 (*45*, *46*) to estimate a species tree, using default settings and 100 bootstrap replicates to calculate node support.

To estimate a time-calibrated tree, we used a penalized likelihood method implemented in treePL (*229*, *230*) and five fossil calibration points (Supplementary Table 3). We first carried out an analysis to determine the optimal optimization parameters for treePL, and subsequently ran a second analysis incorporating these optimized settings. Fossil calibrations were applied to constrain minimum divergence times at relevant nodes.

### Transcriptome meta-assembly and annotation support

Using the transcriptomics data of *Desmodus* and *Diphylla*, we generated RNA-seq meta-assemblies using the xasm workflow (v0.0.20) (https://github.com/hillerlab/xasm). Sequencing reads were converted to BINSEQ format (*231*) and quality controlled using bqc (v0.0.2). Reads were decontaminated with deacon (v0.15.0) (*232*) using default parameters and bacterial and viral genomes. Quality-controlled and decontaminated reads were aligned to the corresponding species genome using a custom two-pass STAR workflow (v2.7.11b) (*233*). During the first pass, all samples were aligned using TOGA2-derived splice junctions as alignment hints. Splice junctions detected across all samples were subsequently aggregated. Additional splice junctions supported by more than five reads, detected in more than two independent samples, and found in introns ≥50 bp in length were retained as a high-confidence junction set. Each sample was then realigned in a second STAR pass using this high-confidence junction set provided through the --sjdbFileChrStartEnd parameter. Both passes used the following STAR parameters: --outSAMtype BAM SortedByCoordinate --outSAMattributes NH HI AS nM NM MD ch -- readFilesCommand zcat --outSAMstrandField intronMotif --outSAMunmapped None -- outFilterMultimapNmax 20 --alignSJDBoverhangMin 1 --peOverlapNbasesMin 5 -- outBAMcompression 10 --limitSjdbInsertNsj 2500000 --outWigType bedGraph --outWigStrand Unstranded --outWigNorm RPM. Dataset-wide genome coverage tracks generated by STAR were processed using bigtools (v0.5.6) (*234*). Coverage at each genomic position was summarized as the median RPM-normalized coverage across samples.

Transcript assemblies were generated with aletsch (v1.1.3) (*235*) using default parameters. To obtain a meta-assembled transcriptome, we integrated the resulting sample-specific transcriptomes with Beaver (v1.0.0) (*236*) using default parameters.

We used the *Desmodus* transcriptome meta-assembly to assess how many TOGA-annotated exons in intact transcripts are supported by transcriptomic evidence. An exon was considered fully supported if both annotated splice sites matched an assembled transcript exon exactly. For terminal exons, only the single annotated splice site was required to match. Because not all genes are expressed in the tissues we profiled, we restricted the analysis to transcripts for which at least one annotated exon overlapped an assembled transcript. Despite this filter, the benchmark provides a lower-bound estimate of annotation support, because lowly expressed genes may lack assembled full-length transcripts, causing correctly annotated exons to be counted as unsupported. For example, whereas *MLN* was lost independently in *Diaemus* and *Diphylla*, *Desmodus* retains an intact gene, but its transcripts cover only the two C-terminal exons, leaving the two N-terminal exons without transcriptomic support and thus counted as unsupported in our benchmark.

Overall, 96.77% of TOGA-annotated exons in the analyzed transcripts are fully supported by transcriptomic data. Considering only the intact genes discussed in the main text, 98.04% of the annotated exons are fully supported.

### Gene losses

We used TOGA’s gene classification to identify inactivated (lost) genes based on the presence of gene-inactivating mutations (frameshifts, premature stop codons, splice site disruptions, and deletions of entire coding exons). We extracted all genes containing at least one inactivating mutation within the middle 80% of the coding region (TOGA status L=loss or UL=uncertain loss) and retained those showing evidence of gene loss in at least one vampire bat species but in no more than five non-vampire bat species. We allowed apparent gene losses in non-vampire bats to avoid false negatives caused by low genome assembly quality. To ensure accuracy, we manually verified each candidate by examining inactivating mutations, codon alignment plots, genomic context, and alignment chains (Supplementary Data 1).

### Validating inactivating mutations and gene losses

To rule out the possibility that the reported gene losses are artifacts, we performed several validation steps. First, for all 23 lost genes presented in the manuscript (Supplementary Table 19), we manually inspected the human to vampire bat genome alignment chains in the genome browser to ensure that they correspond to the orthologous locus. We also verified that no co-orthologous locus containing an intact copy of the gene exists.

Second, we used TOGA2 (*237*) with updated input annotations and four reference species (human, mouse, cow, and elephant) to independently assess the presence of inactivating mutations and gene losses. TOGA2 has increased power to identify evolutionary splice site shifts and changes in exon-intron structure, and confirmed the loss of each gene with each reference species. The only technical complication concerns *CTRB2*, which is genuinely lost in all three vampire bat species. In mammals, *CTRB2* is located in reverse orientation upstream of the conserved and intact paralog *CTRB1*. At this locus, a second alignment chain aligns *CTRB2* to *CTRB1* and *CTRB1* to *CTRB2* in reverse orientation. TOGA2 therefore identifies this second chain as a potential *CTRB2* co-ortholog, albeit with a lower probability (0.66), resulting in a many-to-many orthology relationship in which *CTRB2* is assigned two orthologs: the true *CTRB2* locus, which is correctly classified as lost, and *CTRB1*, which is incorrectly classified as a functional *CTRB2* ortholog. Importantly, manual inspection of the locus shows that the conserved *ZFP1–CTRB2–CTRB1–BCAR1* gene order is retained in all three vampire bats. Moreover, TOGA2 identifies the true *CTRB2* locus at the correct position and detects the same inactivating mutations that led to its classification as lost. Thus, despite the additional spurious orthology relationship, the *CTRB2* gene loss is unambiguous.

Third, to rule out the possibility that base errors in the genome assemblies were incorrectly interpreted as inactivating mutations, we manually validated all 193 identified inactivating mutations by inspecting the HiFi reads. We mapped the HiFi reads to the respective assemblies using backmap (v0.5; https://github.com/schellt/backmap) with default parameters, which is a wrapper for minimap2 and samtools. None of the 193 inactivating mutations was an assembly base error. Of the 193 mutations (63 in *Desmodus*, 46 in *Diaemus*, and 84 in *Diphylla*), 179 (92.7%) were homozygous. The remaining 14 (7.3%) mutations were polymorphic, having ∼50% of the HiFi reads supporting the non-inactivating allele, which likely reflects ongoing gene erosion. All polymorphic mutations are indicated in the Supplementary Figure legends.

Fourth, we used our gene expression data and the above-described transcript meta-assembly to assess whether remnants of the lost genes in *Desmodus* and *Diphylla* were still transcribed. For intron-containing genes, we searched for spliced RNA-seq reads as evidence of genuine transcription and transcript processing at the locus. For single-exon genes, we just searched for RNA-seq reads overlapping the gene remnants. In 17 cases, we found no evidence of spliced RNA-seq reads or assembled transcripts (Supplementary Table 19). The remaining genes generally showed low levels of transcription, but some of the exon-exon junctions were covered by spliced RNA-seq reads, and transcripts partially matching the exon-intron structures annotated by TOGA2 could be assembled. Importantly, for all genes for which we found any evidence of transcription, we manually verified that the RNA-seq reads overlapping TOGA2-annotated exons supported the corresponding inactivating mutations identified in the genome assembly, as illustrated in Supplementary Figure 31. Thus, even where gene remnants are transcribed, the resulting transcripts can no longer encode functional proteins.

Together, these independent validation steps show that the reported gene losses represent genuine gene losses rather than assembly, alignment, or annotation artifacts. Details of the validation are provided in Supplementary Table 19.

### Positive selection using aBSREL

We used the aBSREL method in the HyPhy v2.5.51 package (*47*) to identify genes under positive selection in the vampire bats. This method implements a sensitive branch-site model to detect phylogenetic branches under positive selection. We used the same codon alignments of 15,481 orthologous genes described in the phylogenomics section above, and ran aBSREL with default parameters, except setting the environmental parameter ENV=’TOLERATE_NUMERICAL_ERRORS=1’. In addition, we applied the MEME method in HyPhy with default parameters to detect codons under episodic positive selection.

We extracted all genes with a significant positive selection signal (corrected P-value < 0.05) on any of the five branches in the vampire bat lineage. We manually curated the list of candidate genes to ensure accuracy (Supplementary Data 1). Specifically, we excluded genes that contain inactivating mutations in vampire bats. We also excluded genes where the selection signal appeared to come from problematic alignment regions (e.g. caused by fast-evolving protein regions or potentially mis-annotated exons), as such alignment regions can create spurious signals in aBSREL analyses.

### Relaxed and intensified selection using RELAX

We used the RELAX method (*48*) in the HyPhy v2.5.51 package to identify genes evolving under relaxed or intensified selection in the vampire bat lineage. Like aBSREL, RELAX analyzes codon alignments of orthologous genes, but it compares two user-defined sets of branches (a test and a reference branch set) to determine whether selection to preserve the encoded protein sequence is relaxed or intensified in the test branches relative to the reference branches. In our analyses, the five branches in the vampire bat lineage were defined as the test set, and all other branches in the phylogeny were defined as the reference set. We generated codon alignments as described in the phylogenomics section above, with one key difference of including remnant sequences of genes classified as lost in the codon alignments. Applying the same filters yielded 16,094 codon alignments, a higher number than in the phylogenomics and aBSREL analysis.

For each gene, RELAX was run with parameters “CPU=1 --starting-points 100 --grid-size 2000 --models Minimal”. We obtained the k value to infer the direction of selection, with k < 1 indicating relaxation and k > 1 indicating intensification, and the P-values, which we corrected for multiple testing over the genes using the Benjamini-Hochberg method. Genes with corrected P-values < 0.05 for relaxation or intensification were retained. Because RELAX results can exhibit stochastic variation, we repeated the analysis ten times and merged the resulting gene lists to generate our final set. We then manually curated this gene list by examining the codon alignment visualizations (Supplementary Data 1). As for aBSREL, we removed genes with problematic alignment regions. We also removed genes lost in nearly all species. Finally, we excluded genes that are “hyperconserved” across all species and contain only very few mutations overall, because their alignments provide limited information for reliable inference of relaxation or intensification.

### Differential expression analysis

RNA-seq reads were quality-checked using FastQC v0.11.9 (https://www.bioinformatics.babraham.ac.uk/projects/fastqc/) and MultiQC v1.12 (*238*). Read pairs were filtered using Trimmomatic v0.39 (*239*) setting the following parameters: “ILLUMINACLIP:TruSeq3-PE.fa:2:30:10 LEADING:3 TRAILING:3 SLIDINGWINDOW:4:15 MINLEN:36”.

To quantify gene-level expression, we used kb-python (v0.29.5) (*49*), a wrapper that leverages the pseudoalignment algorithm implemented in kallisto (*240*) to map sequencing reads directly to transcript sequences rather than performing genome-based alignment. We used TOGA coding sequence annotations as input transcripts. To reduce spurious read assignments, we included the genome sequence with hard-masked coding regions as decoy. Kb-python outputs two expression measures for each gene: read count and transcripts per million (TPM). Read counts were used for differential gene expression analysis in DESeq2 (below). To visualize gene expression across tissues, we normalized TPM values with the median-of-ratios method in DESeq2.

To identify genes differentially expressed in different organs between vampire and non-vampire bats, we used DESeq2 v1.46.0 (*241*). We excluded genes with 1:many or many:many orthology relationships according to TOGA, and excluded genes with a median TPM < 0.5 in both vampire and non-vampire bats. Genes with a log2 fold change > 1.5 and adjusted P-value < 0.01 were considered differentially expressed genes.

### Functional enrichment analysis

Gene lists obtained from our comparative genomic and differential expression analyses were subjected to functional enrichment analysis using Metascape (database update: 2025-07-01) (*242*). The following ontological categories were included in the analysis: GO Biological Process, GO Molecular Function, GO Cellular Component, KEGG Pathways, WikiPathways, Canonical Pathways, Reactome, Cell Type Signatures, DisGeNET and PaGenBase. We only considered enrichments with an adjusted P-value < 0.05. The complete enrichment results are provided in Supplementary Tables 11-14 and 17-18.

### Insulin and insulin receptor analysis

To analyze the prevalence of mutations in *INS* and *INSR* in an extended set of mammals, we used TOGA2 (https://github.com/hillerlab/TOGA2) to identify 1:1 orthologs of both genes in available mammalian genomes, which includes bat species that represent all 21 currently recognized bat families (*243*). A multiple alignment was generated with Prank (*244*).

To examine the location of INS mutations in an INS-INSR structural model, we used the crystal structure of the extracellular domain of the insulin receptor bound to insulin (PDB 6PXV), refined using the Protein Preparation Wizard in the Maestro suite (*245*, *246*). Missing side chains and loops were built using the Prime module.

### Estimation of amino acid frequencies in cattle and chicken blood

We estimated whole-blood amino acid frequencies from the amino acid compositions of the two major blood proteins: hemoglobin (HBA1, HBA2 and HBB for cattle; HBAA, HBAD, and HBB for chicken) and albumin. For hemoglobins, the starting amino acid, methionine, was excluded as it is cleaved from the mature globin chain. For albumins, the signal peptide and propeptide were excluded to reflect the mature protein sequences (the N-terminal 24/26 amino acids in cattle/chicken (*247*)). For each species, the amino acid frequencies of these proteins were combined according to their relative abundances in blood. We used the hemoglobin/albumin ratios (7.20 for cattle; 17.33 for chicken) reported by Scanes et al. (2022) (*248*). For chicken, we took the relative abundance of the two adult hemoglobin isoforms, HbA and HbD, into account, using the HbA/HbD ratio of 2.33 reported by Baumann et al. (1984) (*249*), and calculated their contributions based on the different α-globin chain and the shared β-globin chain sequence. The resulting weighted amino acid frequencies provide an estimate of the overall amino acid composition of blood proteins in cattle and chicken and were plotted in the main text figure.

### Visualization

Schematics in the main figures were created using BioRender.com. Sequence logos for INS and INSR were generated with the R package ggseqlogo (https://cran.r-project.org/web/packages/ggseqlogo/index.html).

### Histological analysis of stomach

We used histological methods to investigate three aspects of vampire bat stomach: connective tissue content, mucosa thickness and mucus secretion profile. For this purpose, we captured four *Desmodus,* two *Diphylla*, five *Artibeus lituratus* and four *Myotis nigricans* individuals. Animals were euthanized by decapitation, and the digestive tract was dissected. Fragments of the fundic stomach were washed in saline, then fixed in buffered formalin for 24 h, and stored in 70% ethanol until processing. Fragments were dehydrated in alcoholic solutions of increasing concentration (80/90/100%; 30 min each), immersed in glycol methacrylate resin overnight followed by 2 h in fresh resin. Fragments were subsequently embedded using plastic molds and resin with added polymerizer (hardener). The resulting blocks were sectioned using a rotary microtome and glass blades. The sections obtained were 3 μm thick, with ten semi-serial sections per histological slide (30 μm between sections). Two slides were prepared for each fragment; one was stained with Toluidine blue, and the other with Periodic acid-Schiff conjugated with Alcian blue (PAS-AB). The slides were dried and mounted with coverslips using Entellan resin. Photomicrographs were acquired using a microscope with a camera and image capture system.

Images from Toluidine blue-stained slides were used to quantify the connective tissue content and mucosa thickness, as illustrated in Supplementary Figure 5. Images were annotated for connective tissue, mucosa and muscle using QuPath v0.6.0 (*250*). Connective tissue content was measured both as (i) the percentage of the stomach wall area, and (ii) connective tissue thickness as a percentage of the muscular region (Supplementary Figure 5). Differences between vampire bats and control species were tested using two-tailed Welch’s t-tests. Images from PAS-AB-stained slides were used to assess the mucus secretion profile, with vacuoles showing magenta and blue colors indicating neutral and acidic mucus, respectively.

### Stomach pH

We measured the stomach pH in *Desmodus* (n=4), *Diphylla* (n=5), and *Artibeus lituratus* (n=3). Animals were euthanized by decapitation. For each animal, stomach contents were removed with a small spatula and transferred to an Eppendorf tube, which was then filled with water to the 2 mL mark. The material was further diluted to 20 mL in a test tube for pH measurement using a pH meter. The pH meter recordings are provided in Supplementary Table 21 and shown in the main text figure.

### Gastrointestinal motility

We examined and video-recorded gastrointestinal motility in *Desmodus* (n = 3), *Diphylla* (n = 2), *Artibeus lituratus* (n = 2), *Phyllostomus discolor* (n = 3), and *Mus musculus* (n = 3). Animals were euthanized by decapitation or overdose of isoflurane, and then dissected to expose the abdominal cavity. Gastrointestinal motility was recorded before and after administration of neostigmine (0.5 mg/mL), a cholinergic stimulant of gastrointestinal motility in mammals. Videos of the recordings are provided in Supplementary Videos 1–5.

### Ferritin processed pseudogene copies

Motivated by a previous study that identified an expansion of ferritin genes in the common vampire bat genome (*25*), we reanalyzed ferritin genes in our extended set of genomes. As shown in Supplementary Figures 27-29, we analyzed *FTL* and *FTH1* orthologs detected by TOGA and inspected the alignment chains in the *FTL* and *FTH1* locus, which revealed a single *FTH1* ortholog, an ancient *FTL* duplication, and for all analyzed bats a large number of processed pseudogene copies. To determine whether there is a vampire bat specific expansion of ferritin processed pseudogene copies, we compared the three vampire bats with four other species that have long read-based genomes, representing non-vampire phyllostomid bats (*Micronycteris megalotis*, *Carollia perspicillata*, *Glossophaga soricina*) and the outgroup *Mormoopidae* family (*Pteronotus mesoamericanus*). Since RepeatModeler may detect processed pseudogenes as an unknown repeat family if they have several hundred copies in an assembly, we removed any soft-masking corresponding to repeat families with sequence similarity to *FTL* or *FTH1* before computed alignment chains. To systematically compare the number of processed pseudogene copies, we leveraged a new functionality of TOGA2 (https://github.com/hillerlab/TOGA2) to detect processed pseudogenes and determine whether they have an intact open reading frame. To detect all processed pseudogenes, we lowered the minimum chain score from 15,000 to 5,000 for these TOGA2 runs. The number of detected processed pseudogenes and how many have intact reading frames are listed in Supplementary Table 22.

## Acknowledgment

We thank Leandro Costa do Nascimento and Maria Carolina Svidnicki for RNA-sequencing, Matheus Canal, Monika Fischer, Sasha Newar, Emilia NovilloSuarez and Tom Jenks for helping with experiments, and the Genome Technology Center (RGTC) at Radboudumc for the use of the Sequencing Core Facility (Nijmegen, The Netherlands), which provided the PacBio SMRT sequencing service on the Sequel IIe platform, the Environment Services and Support (ESS) in Suriname. We thank Brock Fenton, Merlin Tuttle, José G. Martínez-Fonseca and Jon Flanders for permission to use bat photographs. For support in the Colombia 2019 expedition, we thank Andrés Cuervo, Mailyn Gonzalez, Andrés Julián Lozano Florez, Darwin M. Morales Martinez, Paola Pulido Santacruz, Danny Rojas, and Laurel Yohe. We acknowledge support by the Sector Plan Pharmaceutical Sciences, implemented in the overarching Sector Plan Beta II and put into action by the Ministry of Education, Culture, and Science (OCW) of The Netherlands.

This work was supported by the German Research Foundation (grants HI1423/5-1 and HI1423/6-1 to MH, and HI2214/1-1 to LH), the LOEWE-Centre for Translational Biodiversity Genomics (TBG) funded by the Hessen State Ministry of Higher Education, Research and the Arts (HMWK) (LOEWE/1/10/519/03/03.001(0014)/52), the Max Planck Society, and the European Research Council (ERC-2023-SyG, 101118919). LMD was supported by National Science Foundation Grants NSF RoL: FELS: EAGER 1838273 and NSF-IOS 2032063 and 2031906, and a Colciencias Seed Grant. BKL received funding from Environmental Services & Support (ESS) for fieldwork in Suriname, the Royal Ontario Museum Department of Natural History for fieldwork in Guyana, and the Ecuambiente Consulting Group for fieldwork in Ecuador. SCV was supported by a UKRI Future Leaders Fellowship (MR/T021985/1; UKRI2048) and ERC Consolidator Grant (101001702; BATSPEAK). Genome sequencing of non-vampire bats was performed by the LongRead Project of the Max-Planck Institute of Molecular Cell Biology and Genetics (MPI-CBG) as part of the DcGC Dresden-concept Genome Center and was supported by the Next Generation Sequencing Competence Network (DFG project 423957469).

## Data availability

Genomic and transcriptomic data and the genome assemblies of *Desmodus rotundus* (PRJNA1557061, PRJNA789159), *Diaemus youngii* (PRJNA1556840, PRJNA1256700), *Diphylla ecaudata* (PRJNA1556832, PRJNA1256387), *Micronycteris megalotis* (PRJNA1557323, PRJNA1263771), *Carollia perspicillata* (PRJNA1265070), and *Glossophaga soricina* (PRJNA1557329, PRJNA1265439), and transcriptomic data for *Artibeus obscurus* (PRJNA1399465) and *Leptonycteris yerbabuenae* (PRJEB1149427, PRJEB107355) are available on NCBI under the listed BioProject IDs.

## Code availability

No new code was generated for this study.

